# Heterotrophic flagellates and centrohelid heliozoans from marine waters of Curacao, the Netherlands Antilles

**DOI:** 10.1101/2020.08.10.243865

**Authors:** Kristina I. Prokina, Patrick J. Keeling, Denis V. Tikhonenkov

## Abstract

Recent progress in understanding the early evolution of eukaryotes was tied to morphological identification of flagellates and heliozoans in the natural samples, isolation of their cultures and genomic and ultrastructural investigations. These protists are the smallest and least studied microbial eukaryotes but play an important role in functioning of microbial food webs. Using light and electron microscopy, we have studied the diversity of heterotrophic flagellates and centrohelid heliozoans from marine waters of Curacao (The Netherlands Antilles), and provide micrographs and morphological descriptions of observed species. Among 86 flagellates and 3 centrohelids encountered in this survey, five heterotrophic flagellates and one Centrohelid heliozoan were not identified even to the genus. Some flagellate protists have a unique morphology, and may represent undescribed lineages of eukaryotes of high taxonomic rank. The vast majority (89%) of identified flagellates are characterized by wide geographical distribution and had been reported previously from all hemispheres and various climatic regions. More than half of the species were previously observed not only from marine, but also from freshwater habitats. The parameters of the species accumulation curve indicate that our species list obtained for the Curacao study sites is far from complete, and each new sample should yield new species.

## INTRODUCTION

Heterotrophic flagellates and centrohelid heliozoans are the smallest and least studied groups of protists both at the morphological and molecular levels. Heterotrophic flagellates – the collective name for an extremely diverse “hodgepodge” of polyphyletic, colorless protists moving or feeding with flagella at least in one stage of their life cycle (Patterson and Larsen 1991) – are characterized by significantly variety of metabolism and ecology. At the same time centrohelid heliozoans (Centroplasthelida Febvre-Chevalier et Febvre, 1984) is a monophyletic group of predatory protists, related to haptophyte algae within the supergroup Haptista Cavalier-Smith, 2003 (Burki et al. 2016). Centrohelids are characterized by the presence of surface cytoskeletal structures – siliceous scales or organic spicules, having an important diagnostic value.

Heterotrophic flagellates and heliozoans are widespread in different types of freshwater and marine biotopes and play an important role in functioning of the microbial food webs, that provide effective pathways for the transformation of matter and energy in aquatic ecosystems (Arndt et al. 2000; Domaizon et al. 2003; Kiss et al. 2009). Beyond that, the contribution of the eukaryovorous centrohelids and flagellates to the trophic structure and functioning of aquatic ecosystems is also probably very important, but not fully understood.

Faunistic investigations of heterotrophic flagellates and heliozoans supported with morphological descriptions were especially common in protistological literature in the 90s – early 2000s (Larsen and Patterson 1990; Lee and Patterson 2000; Lee et al. 2003; Mikrjukov 2001; Patterson and Simpson 1996; Patterson et al. 1993; Tong 1997b, c; Vørs 1992). Such studies have fallen from favour, and their scientific importance is often undeservedly disregarded, having been largely replaced by molecular surveys from bulk microbial community DNA (Geisen et al., 2019; Pawlowski et al., 2016). However, several recent advances in the study of eukaryotic evolution have been specifically tied to morphological identification of flagellates and heliozoans in the natural samples, their isolation inculture, and genomic and ultrastructural investigations from those cultures. Morphological recognition and identification (sometimes raw and approximate) of these protists in samples is precisely based on such “classical” studies, and would not have been possible without them. For example, recent recognition, isolation and investigation of previously understudied or unknown heterotrophic flagellates were essential in addressing major evolutionary problems, such as the origins of photosynthesis and parasitism and the trajectory of plastid spread (Gawryluk et al. 2019; Janouškovec et al. 2015; Tikhonenkov et al. 2020b), origin of multicellular animals (Hehenberger et al. 2017; Tikhonenkov et al. 2020a), evolution of mitochondrial genomes, rooting of the tree of eukaryotes and clarification of their relationships (Janouškovec et al. 2017; Lax et al. 2018; Strassert et al. 2019). Identification and establishing of clonal cultures of centrohelids were crucial for transcriptomic research and untangling the early diversification of eukaryotes (Burki et al. 2016).

Descriptions of many flagellates and heliozoans were made in the late XIX and early XX centuries using imperfect light microscopy: many of these species lack type material, ultrastructural, and molecular data, and the taxonomy of many species and groups is in need of revision (Lee et al. 2003; Schoenle et al. 2020). Currently, the described diversity of these protists represents only a small fraction of their total species richness in nature (Cavalier-Smith and von der Heyden 2007; Corliss 2002). Metagenomic and metabarcoding sequencing of environmental samples has revealed several lineages representing high levels of hidden diversity: e.g. ribogroups MALV, MAST, MAOP, MAFO, deep-sea pelagic diplonemids (DSPD), or eupelagonemids (del Campo and Ruiz-Trillo 2013; del Campo et al. 2015; Guillou et al. 2008; Massana and Pedrós-Alió 2008; Okamoto et al. 2019; de Vargas et al. 2015), as well as great variety of small new phylogenetic lineages associated with almost all large eukaryotic groups (del Campo et al. 2016; Keeling and del Campo 2017). New species of flagellates and centrohelids are constantly being discovered, which is indicative of a poor state of exploration and insufficient sampling.

At the same time, the questions about geographical distribution of protists are subject to lively, but stubbornly unresolved debate (Azovsky et al. 2016). Are protists species widespread around the globe, or is their distribution, like those of macroorganisms, more governed by the rules of historical biogeography? There are two alternative points of view on this issue: the Ubiquity Model (Fenchel and Finlay 2004; Finlay et al. 1996) and Moderate Endemicity Model (Foissner 2004, 2008). Additional data is needed to resolve this issue, especially in those regions on the planet that remain unexplored.

Here we describe a microscopical investigation of the species diversity of heterotrophic flagellates and centrohelid heliozoans in marine waters of coast of Curacao, which has not specifically been investigated previously. For the Caribbean Sea in general, only seven species of choanoflagellates were known (Thomsen and Østergaard 2019) until recent. We find the diversity of heterotrophic protists on Curacao to be rich and intriguing, including the presence of species potentially important for clarifying previously puzzling evolutionary and ecological questions. For example, the recently described colonial flagellate *Choanoeca flexa*, whose discovery and description from Curacao significantly contributed to the understanding of the origin of polarized cell contractility in animals (Brunet et al. 2019). We describe a survey of light and electron microscopical studies of the diversity of heterotrophic flagellates and centrohelid heliozoans from marine waters of several locations on Curacao, and provide micrographs and morphological descriptions of observed species, as well as discussing their distribution and potential importance.

## MATERIALS AND METHODS

Marine water samples were taken from eight locations around the island of Curacao in April 2018. Samples came from several biotopes (from the surface of corals and sponges, *Sargassum* algae wrings, sand, coral sand, and water column) both nearshore and at depths of 12–25 m (detailed descriptions of sampling points are given in the Table). From each biotope, several replicates were taken, which were subsequently summarized. A total of 52 samples were investigated microscopically. Water temperature in the studied region is relatively constant, about 26–29°C. Samples were placed into 50 ml plastic tubes and transported to the laboratory at 4°C.

Samples were enriched with a suspension of *Pseudomonas fluorescens* Migula, 1895 bacteria and placed in Petri dishes. Samples were kept at 22°C in the dark and observed for 10 days to reveal the cryptic species diversity (Tikhonenkov et al. 2008; Vørs 1992). A culture of the kinetoplastid flagellate, *Procryptobia sorokini* (Zhukov, 1975) Frolov et al., 2001, was used as a food source for predatory heliozoans and flagellates.

An AxioScope A1 light microscope (Carl Zeiss, Germany) with DIC and phase contrast and water immersion objectives (total magnification ×1120) was used for observations of living cells. Electron microscope preparations were carried out according to described methods (Mikrjukov 2002; Moestrup and Thomsen 1980) and observed in a JEM-1011 (Jeol, Japan) transmission electron microscope and a JSM–6510 LV (Joel, Japan) scanning electron microscope.

The dendrogram showing the similarity of types of biotopes by flagellates’ species composition was drawn on the basis of the Dice similarity index using the paired group algorithm in the PAST software package (Hammer et al. 2001).

The analysis of geographic distribution of heterotrophic flagellates was based on “morphospecies” concept (Fenchel and Finlay 2006; Finlay et al. 1996) and carried out using the previously published database (Azovsky et al., 2016, 2020) and available literature sources. Only morphology-based (or morphology-confirmed) data were considered.

## RESULTS

Eighty-six species and forms of heterotrophic flagellates and three species of centrohelid heliozoans were observed, all of them are listed below systematically. A system of asterisks was used to identify higher levels of taxonomic ranks (Order, sub-order, Super Family, Family, etc.) per Adl et al. (2019). Morphological characteristics of most unusual and rare species of flagellates and all centrohelids are gived. For each species, sample points are presented in accordance with the Table. Presumably cosmopolitan morphospecies, indicated with ^“c”^, was previously recorded from all hemispheres (North, South, West, and East) and from equatorial, tropical or subtropical, temperate, and polar regions.

AMORPHEA Adl et al., 2012

Obazoa Brown et al., 2013

*Apusomonadida Karpov et Mylnikov, 1989

*Amastigomonas debruynei* De Saedeleer, 1931^c^ – observed in sample points 2e, 3a, 3c, 8a.

*Podomonas griebenis* (Mylnikov, 1999) Cavalier-Smith in Cavalier-Smith et Chao, 2010^c^ –observed in sample points 3a, 5b, 8b.

*Thecamonas mutabilis* (Griessmann, 1913) Larsen et Patterson, 1990^c^ – 2a, 6

***Thecamonas* sp.** (Fig. 1a–c) – observed in sample point 7.

**Fig. 1.**
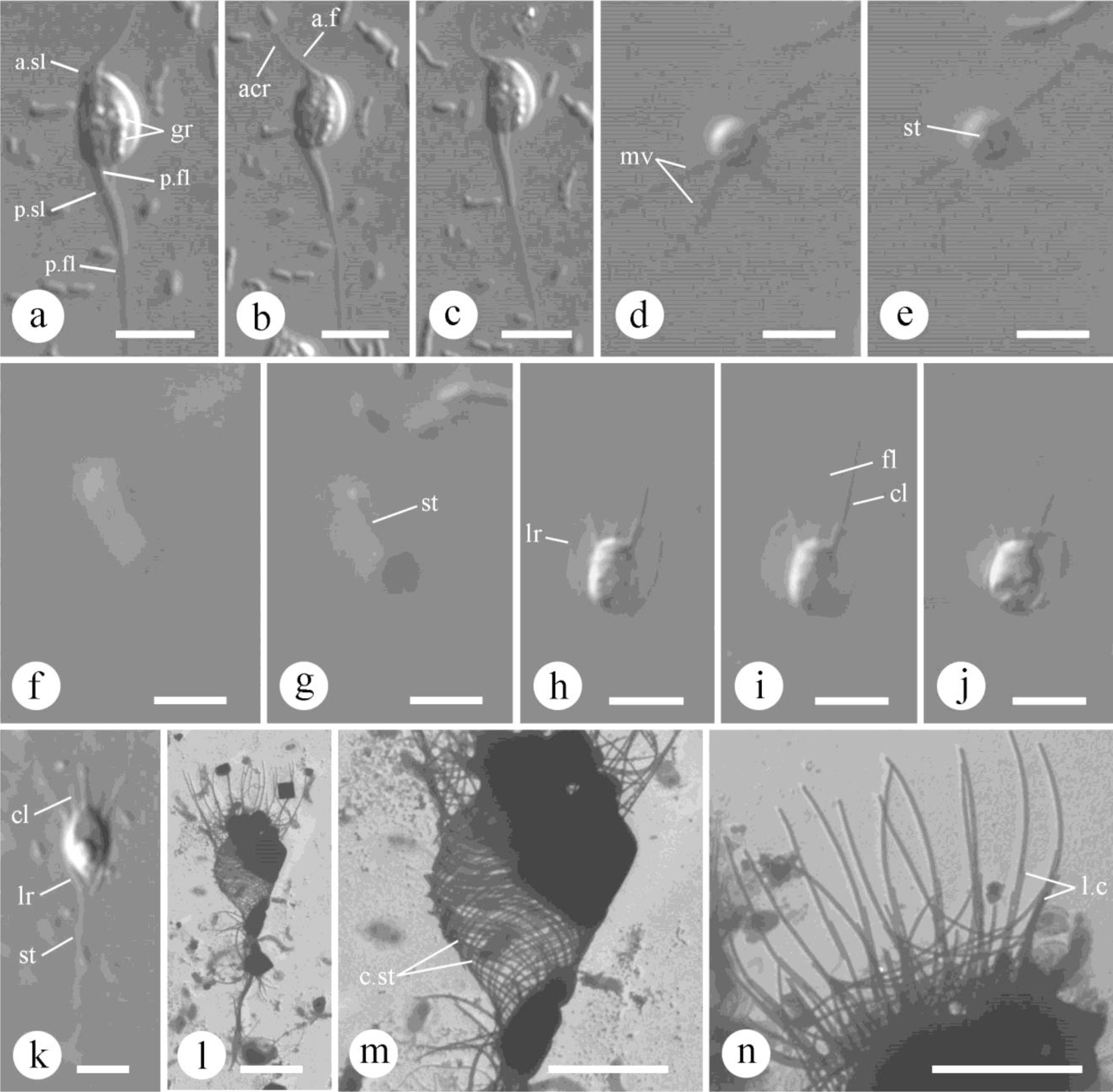
Morphology of observed heterotrophic flagellates (a–k – DIC, l–n – TEM): a–c – *Thecamonas* sp.; d–g – *Ministeria vibrans* (d, e – non-moving cell, f–g – moving cell); h–j – *Salpingoeca pyxidium*; k–n – *Acanthoeca spectabilis* (k – living cell, l – whole lorica; m –posterior chamber of the lorica; n – anterior chamber of the lorica). Abbreviations: acr – acronema; a.f – anterior flagellum; cl – collar; c.st – costal strips; fl – flagellum; gr – granules; lr – lorica; l.c – longitudinal costa; mv – microvilli; p.f – posterior flagellum; ps – pseudopodia; sl – sleeve; st –stalk. Scale bars: 5 µm.

### Observations

Cells are oval or reniform in shape, dorso-ventrally flattened, 5–6 μm in length, 3.5–4.0 μm in width. Anterior flagellum is about the cell length, its basal part passes in the sleeve, distal part is acronematic. Posterior flagellum is 4 times longer than the cell, along half-length passes in the sleeve. Numerous granules are visible inside the cell. The cells move by gliding, with anterior flagellum beating with a flicking motion. Observation based on description of six cells in LM.

### Remarks

Cavalier-Smith and Chao (2010) divided marine *Amastigomonas*-like species into four genera: *Manchomonas*, *Multimonas*, *Podomonas*, and *Thecamonas*. The last genus was introduced earlier by Larsen and Patterson (1990) for biflagellate *Amastigomonas*-like protists whose anterior flagellum and dorsal cell body surface are covered by a pliable organic theca. We assigned species to the genus *Thecamonas* due to the presence of moderately rigid cell covering (probably, theca). This ogranism differs from *Manchomonas* species by the presence of acronematic flagella; one long posterior flagellum that extends beyond the ventral groove, and a small and weakly noticeable triangular sleeve. We didn’t observe multicellular plasmodia with incomplete cytokinesis, as seen in *Multimonas* species, in Petri dishes over the course of 10 days observation. Observed cells differed from *Podomonas* species by the absence of multiform finger-like, lamellar or reticulose pseudopodia. Ultrastructural and molecular data are needed for confirmation of position of this species within the genus *Thecamonas*.

Observed species differ from all known *Amastigomonas*-like species, except *Podomonas* (=*Amastigomonas*) *klosteris* (Arndt et Mylnikov in Mylnikov, 1999) Cavalier-Smith in Cavalier-Smith et Chao, 2010, because of the presence of long posterior sleeve surrounding the posterior flagellum. However, *Podomonas klosteris* is characterized by larger cell body (length 12–20 μm) with posterior sleeve that is shorter relative to the cell (about one quarter of the cell body length), and presence of pseudopodia emerged from ventral groove (Mylnikov 1999).

**\***Opisthokonta Cavalier-Smith, 1987

**Holozoa Lang et al., 2002

***Filasterea Shalchian-Tabrizi et al., 2008

***Ministeria vibrans*** Tong, 1997 ^c^ (Fig. 1d–g) – observed in sample points 6, 8b. <u>Observations</u>: Spherical cell body is 2.5–3.0 μm in diameter, with 10–15 radial thin microvilli (arms) 4–5 μm in length. Cells attach to the substrate with a stalk (Fig. 1g) and make fast pendulum movements with small amplitude (Fig. 1f). Sometimes cells stop moving for a while. Flagellates feed on bacteria, predominantly rod-like, that adhere to the surface of the cell body. Nine cells observed in LM.

### Remarks

Mylnikov and coauthors (Mylnikov et al. 2019) showed that *Ministeria* arms are supported by bundles of axial microfilaments that correspond in structure to the microvilli surrounding the collar of choanoflagellates, and which can be drawn into the cell body. This species differs from another member of the genus, *M. marisola* Patterson et al., 1998, by having a larger number of radial filopodia, as well as attachment to a substrate with a stalk and pendulum movements (Tong et al. 1997b).

### Distribution

Marine waters of UK (Tong 1994), Australia (Lee et al. 2003), Black Sea (Prokina et al. 2018), Baltic Sea (Mylnikov et al. 2019). Fresh waters of European part of Russia, Kostroma Region (unpublished), Hungary (Kiss et al. 2009). Hypersaline inland waters of Southern Chile, Tierra del Fuego (unpublished).

***Choanoflagellata Kent, 1880–1882

****Craspedida Cavalier-Smith, 1997

*****Salpingoecidae Kent, 1880–1882 *sensu* Nitsche et al., 2011

*Salpingoeca infusionum* Kent, 1880–1882^c^ – 1b, 4d *S. minor* Dangeard, 1910^c^ – 5a ***Salpingoeca pyxidium*** Kent, 1880–1882^c^ (Fig. 1h–j) – observed in sample point 4d. <u>Observations</u>: Cells located into organic lorica, that is heart-shaped in outline: narrowed basal part and broad anterior part. Closer to the mouth of the lorica, its edges curved towards the cell and forming a small indentation. Lorica length is 5–6 μm, width at the expanded anterior part is 4.5–5.5 μm. Cells obovoid, 4–5 μm long, 3.0–3.5 μm wide, with slightly expanded basal part and truncated anterior end. Collar length is 1–2 times longer than the cell, located mostly outside the lorica. Single flagellum is slightly longer than the cell. Nucleus located medially. Several food vacuoles located posteriorly. Stalk wasn’t observed, cells attached to the substrate with basal part of the lorica. Three cells observed in LM.

### Remarks

Some authors described the lorica as almost circular in outline (Francé 1897; Lemmermann 1910; Starmach 1968; Tikhonenkov et al. 2008; Zhukov and Karpov 1985), although Kent in the original description (1880) pointed out the heart-shaped morphology of the lorica. The morphology of the cells we observed corresponded to the morphology described by Kent. Sizes of cells are in agreement with previous descriptions, except length of flagellum in observed cells is almost equal to cell length, while many authors noted flagellum length is 2.5 times longer than the cell.

### Distribution

Fresh waters of European part of Russia (Kopylov et al. 2015; Tikhonenkov and Mazei 2007), U.K. (Kent 1880), Hungary (Francé 1897), Poland (Starmach 1968). Black Sea (Tikhonenkov 2006), Pechora Sea (Mazei and Tikhonenkov 2006; Tikhonenkov 2006).*S. ringens* Kent, 1880–1882^c^ – observed in sample points 4d, 5b.

1. *S. ringens* Kent, 1880^c^ – observed in sample point 4d, 5b.
2. *S. tuba* Kent, 1880–1882^c^ – observed in sample point 3b.

****Acanthoecida Cavalier-Smith, 1997

*****Acanthoecidae Norris, 1965 *sensu* Nitsche et al., 2011

***Acanthoeca spectabilis*** Ellis, 1930^c^ (Fig. 1k–n) – observed in sample point 7. <u>Observations</u>: The living cell measured was 8.0×4.5 μm, with rounded anterior end and tapered posterior end. Cells located in conical lorica 12.5–16.0 μm in length. Lorica consist of anterior chamber (Fig. 1n) and posterior chamber (Fig. 1m). Anterior chamber is 4.5–6.5 μm in length, 7.5–10.5 μm in diameter, consist of 15–16 longitudinal costae, each of them includes two rod-like costal strips. Posterior chamber is 8–9 μm in length, 5.5–6.0 μm in diameter, consist of numerous helically twisted costae. Lorica attached to the substrate with a long and thick stalk, 11–20 μm in length, 0.4–0.6 μm in diameter. Base of stalk expanded. Stalk consists of many rod-like costal strips oriented parallel to the longitudinal axis of the stalk and tightly adjacent to each other. At the base of posterior chamber, the elements gradually twist into a spiral. Descriptions based on observation of one living cell in LM and six loricae in TEM.

### Remarks

Morphology of observed lorica corresponds with previous descriptions, however, some authors recorded other number of longitudinal costae: 10–16 (Leadbeater 1972), 13–15 (Leadbeater et al. 2008), 14 costae in original descriptions (Norris 1965). This species differs from the second common species of the genus, *A. brevipoda* Ellis, 1930, by the presence of a stalk.

### Distribution

Marine waters of U.K. (Leadbeater and Morton 1974), Denmark (Thomsen 1973), Norway (Leadbeater 1972), USA (Norris 1965), Australia (Lee 2015; Lee et al. 2003; Tong 1997c; Tong et al. 1998), Taiwan (Hara et al. 1997), Antarctica (Marchant and Perrin 1990), Baltic Sea (Vørs 1992). Hypersaline waters of Australia (Al-Qassab et al. 2002).

*****Stephanoecidae Leadbeater, 2011

***Acanthocorbis camarensis*** Hara in Hara et al., 1996 (Fig. 2a–d) – observed in sample points 2a, 3a, 3b, 6, 7.

**Fig. 2.**
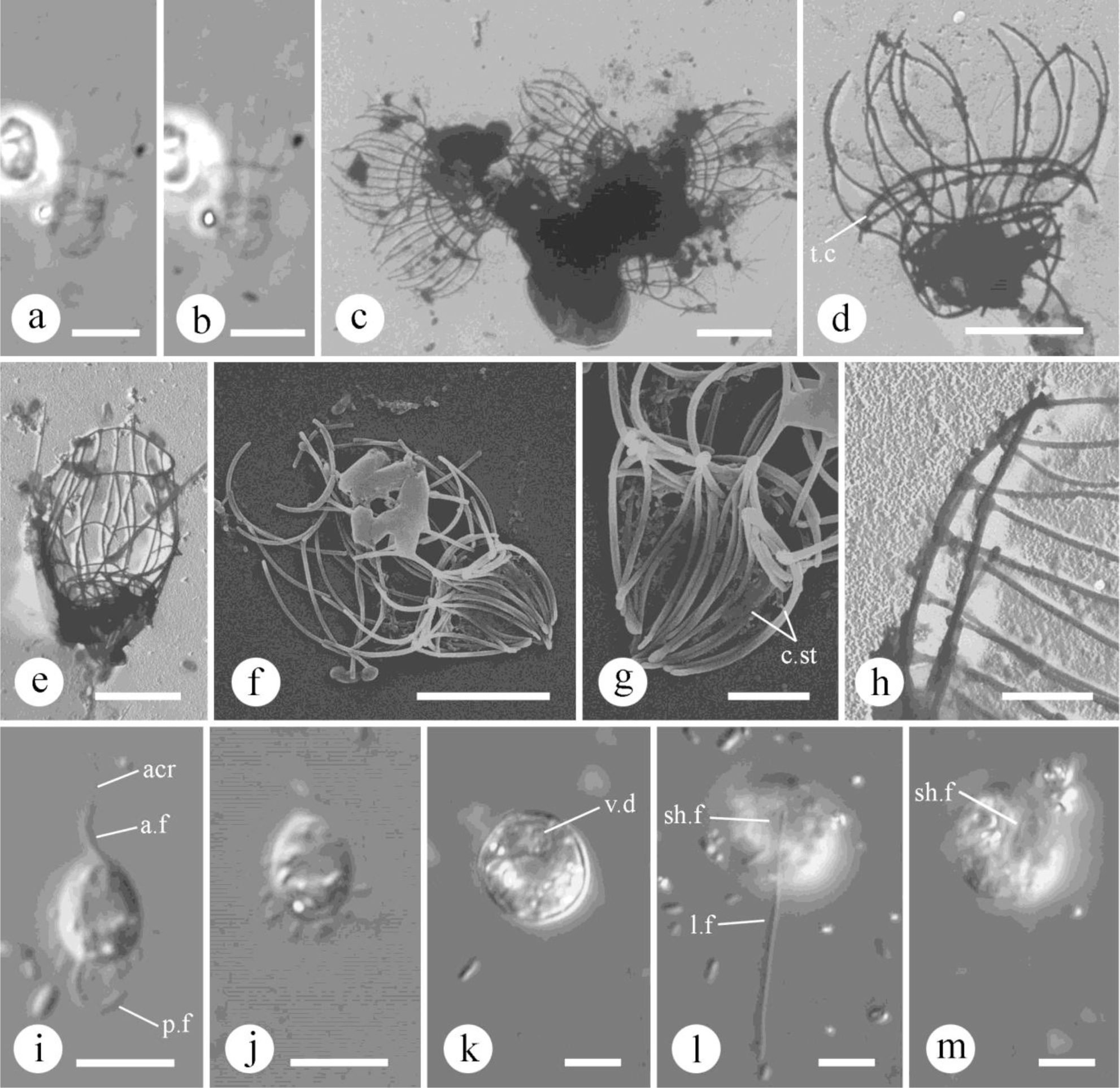
Morphology of observed heterotrophic flagellates (a, b – PhC, c–e, h – TEM, f, g – SEM, i–m – DIC): a–d – *Acanthocorbis camarensis* (a, b – living cell, c, d – loricae); e–h – *Stephanoeca supracostata* (e, f – whole lorica, g – posterior chamber, h – mouth of the lorica); i, j – *Cyranomonas australis* (i – flagella in focus, j – attached particles in focus); k–m – *Clautriavia biflagellatta*. Abbreviations: acr – acronema; a.f – anterior flagellum; c.st – costae strips; l.f – long flagellum; sh.f – short flagellum; t.c – transversal costa; v.d – ventral depression. Scale bars: a–f, i–m – 5 µm; g, h – 2 µm.

### Observations

Lorica is 10.6–14.7 μm in length, consist of two chambers. Anterior chamber is 6.6–9.2 μm in length, 10.3–12.0 μm in diameter, expands in the central part and tapers slightly towards the mouth. Consist of 12–14 longitudinal costae and one double transversal costa (or two transverse costae adjacent). Longitudinal costae consist of three rod-like costal strips, 3.4–4.9 μm in length, 0.15–0.20 μm in diameter. Transversal costa of anterior chamber located closer to the basal part of lorica, consist of many rod-like costal strips, overlapping each other. Posterior chamber bell-shaped, often obscured by protoplast, 3.8–6.3 μm in length, 5.2–5.7 μm in diameter, consist of several transversal and helical costal strips. The number of longitudinal costae of posterior chamber, probably, equal to the number of longitudinal costae of the anterior chamber. Description based on records of one living cell in LM, and eleven loricae in TEM.

### Remarks

Morphology of observed lorica corresponds with previous descriptions, however, Thomsen and Østergaard (2019) indicated up to 18 longitudinal ribs. We recorded only 12–14 (often 14) ribs, which matches the original description, 10–14 longitudinal ribs in Hara et al. (1996). This species close resembles *A. apoda* (Leadbeater, 1972) Hara et Takahashi, 1984 by the presence of a double transverse rib, absence of stalk, as well as the presence of “empty space” between the transverse ribs of anterior and posterior chambers (Leadbeater 1972). However, *A. camarensis* can be distinguished by shorter distance between anterior transverse costa and the lorica waist (less than length of one longitudinal costal strip), and presence of additional helical costal elements in posterior chamber (Thomsen and Østergaard 2019).

### Distribution

Marine water of Hawaii, Thailand, Taiwan (Hara et al. 1996; Thomsen and Østergaard 2019), Australia (Lee et al., 2003).

*Stephanoeca apheles* Thomsen, Buck et Chavez, 1991^c^ – observed in sample points 3b, 6, 7

*S. cupula* (Leadbeater, 1972) Thomsen, 1988^c^ – observed in sample points 3b, 5a.
*S. diplocostata* Ellis, 1930^c^ – observed in sample point 3b.
***Stephanoeca supracostata*** Hara in Hara et al., 1996^c^ (Fig. 2e–h) – observed in sample points 6, 7.

### Observations

Living cells were not observed. Cells surround by basket-like lorica, consist of siliceous costa. Lorica length (according to TEM and SEM whole mount) is 10.5–14.0 μm, diameter in broad middle part is 7.5–8.0 μm. Lorica consist of anterior and posterior chambers. Anterior chamber is 7.0–9.5 μm in length, includes 12–14 longitudinal costae and three transversal costae. Longitudinal costae consist of two rod-like costal strips, connected to each other with a slight overlap. Anterior costal strips have flattened anterior tips, and posterior costal strips have flattened posterior tips. Anterior transversal costae located at the lorica mouth are 5–6 μm in diameter. The second transversal costa is 4.5–5.5 μm in diameter, located on the broadest part of lorica, near the overlap of longitudinal costal strips. The third transversal costa is located at the junction of anterior and posterior chambers. Posterior chamber is 4.0–4.5 μm in length, tapers conically to the basal end, and consist of numerous longitudinal costal strips with both tips flattened. Observation based on description of four loricae in TEM and one lorica in SEM.

### Remarks

Morphology and size of observed cells corresponds with previous descriptions: lorica length is 9–18 μm (Hara et al. 1996; Thomsen and Østergaard 2019); lorica width is 4.5–10 μm (Hara et al. 1996); number of longitudinal costae is 7–20 (Hara et al. 1996; Lee et al. 2003).

The morphology of the observed loricae is very similar to lorica of *S. elegans* (Norris, 1965) Throndsen, 1974. Studied species can be distinguished from *S. elegans* by presence of third middle transversal costa at the broadest lorica’s part, as well as conically tapered posterior chamber. Other species of *Stephanoeca* has more than three transversal costae.

### Distribution

Marine waters of Australia (Lee et al. 2003), Taiwan, Japan (Hara et al. 1996), Thailand, Denmark (Thomsen and Østergaard 2019), U.K. (Tong 1997a).

*Volkanus costatus* (Valkanov, 1970) Özdikimen, 2009^c^ – observed in sample points 3d, 6, 1. 7.

DIAPHORETICKES Adl et al., 2012

SAR Burki et al., 2008

*Alveolata Cavalier-Smith, 1991

**Colpodellida Cavalier-Smith, 1993

***Colpodellaceae Adl et al., 2019

*Colpodella* sp. – observed in sample point 6.

**Colponemida Cavalier-Smith 1993

***Colponemidia Tikhonenkov et al. 2014

*Colponema marisrubri* Mylnikov et Tikhonenkov, 2009 – observed in sample point 3c.

*Rhizaria Cavalier-Smith, 2002

**Cercozoa Cavalier-Smith, 1998

***Thecofilosea Cavalier-Smith, 2003

****Cryomonadida Cavalier-Smith, 1993

*****Protaspidae Cavalier-Smith, 1993

*Protaspa obliqua* (Larsen et Patterson, 1990) Cavalier-Smith in Howe et al., 2011^c^ – observed in sample point 6.

*P. tegere* (Larsen et Patterson, 1990) Cavalier-Smith in Howe et al., 2011^c^ – observed in point 6.

*P. verrucosa* (Larsen et Patterson, 1990) Cavalier-Smith in Howe et al., 2011^c^ – observed in sample point 7.

***Imbricatea Cavalier-Smith, 2011

****Marimonadida Cavalier-Smith et Bass, 2011

***Cyranomonas australis*** Lee, 2002^c^ (Fig. 2i, j) – observed in sample points 2a, 7. <u>Observations</u>: Cells are ovoid in outline, with slightly narrowed anterior end, and dorso-ventrally flattened. Cell length is 4.5–5.5 μm, width is 3.5–4.5 μm. Flagella insert subapically from small ventral depression. Anterior flagellum is 1.0–1.2 times longer than the cell, acronematic, and makes flicking movements from anterior to lateral direction. Posterior flagellum is twice as long as the cell, and trailed behind the cell. Numerous particles attached to the posterior part of the cell (Fig. 2j). Nucleus located anteriorly. Seven cells were observed in LM.

### Remarks

Cell sizes are similar to those described previously (Aydin and Lee 2012; Lee 2002; Tikhonenkov 2009), however, particles attached to the posterior part of the body of the cell were not previously indicated. Also, both non-acronematic flagella were indicated previously (Aydin and Lee 2012; Lee 2002, Tikhonenkov 2009), while we observed acronema on the anterior flagellum.

### Distribution

Marine waters of Turkey (Aydin and Lee 2012), South Korea, Australia (Lee 2002), Red Sea (Tikhonenkov 2009), White Sea ([as Flagellata sp. 1] Tikhonenkov et al. 2006; Tikhonenkov and Mazei 2013).

*Cyranomonas* sp. – observed in sample point 6.

****Variglissida Cavalier-Smith, 2014

***Clautriavia biflagellata*** Chantangsi et Leander, 2009 (Fig. 2k–m) – observed in sample points 3a, 5b, 6, 7.

### Observations

Cells are circular in outline, dorso-ventrally flattened, with slightly concave ventral surface and anterior-medial depression. Cell diameter is 9–13 μm. Two unequal non-acronematic flagella emerge from ventral depression and both directed posteriorly. Long flagellum is 2.0–2.3 times longer than the cell, and trailed behind the cell. Short flagellum is 2.5–3.0 μm in length, immobile, and slightly thinner than the long flagellum. It is weakly visible during normal cell movement (Fig. 2l) and can be noticed when cells rise above the substrate (Fig. 2m).

Pseudopodia were not observed. Four cells were observed in LM.

### Remarks

Morphology of observed cells corresponds with original description (Chantangsi and Leander 2010), at the smaller end of the size range indicated by authors. This species distinguishable from other species of the genus *Clautriavia* by presence of second short flagellum and circular cells in outline. Feeds on algae through the ventral side of the cell (Chantangsi and Leander 2010).

### Distribution

Marine waters of British Columbia, Canada (Chantangsi and Leander 2010).

****Silicofilosea Adl et al., 2005

*****Thaumatomonadida Shirkina, 1987

******Thaumatomonadidae Hollande, 1952

***Thaumatomonas seravini*** Mylnikov et Karpov, 1993^c^ (Fig. 3a–c) – observed in sample point 5b.

**Fig. 3.**
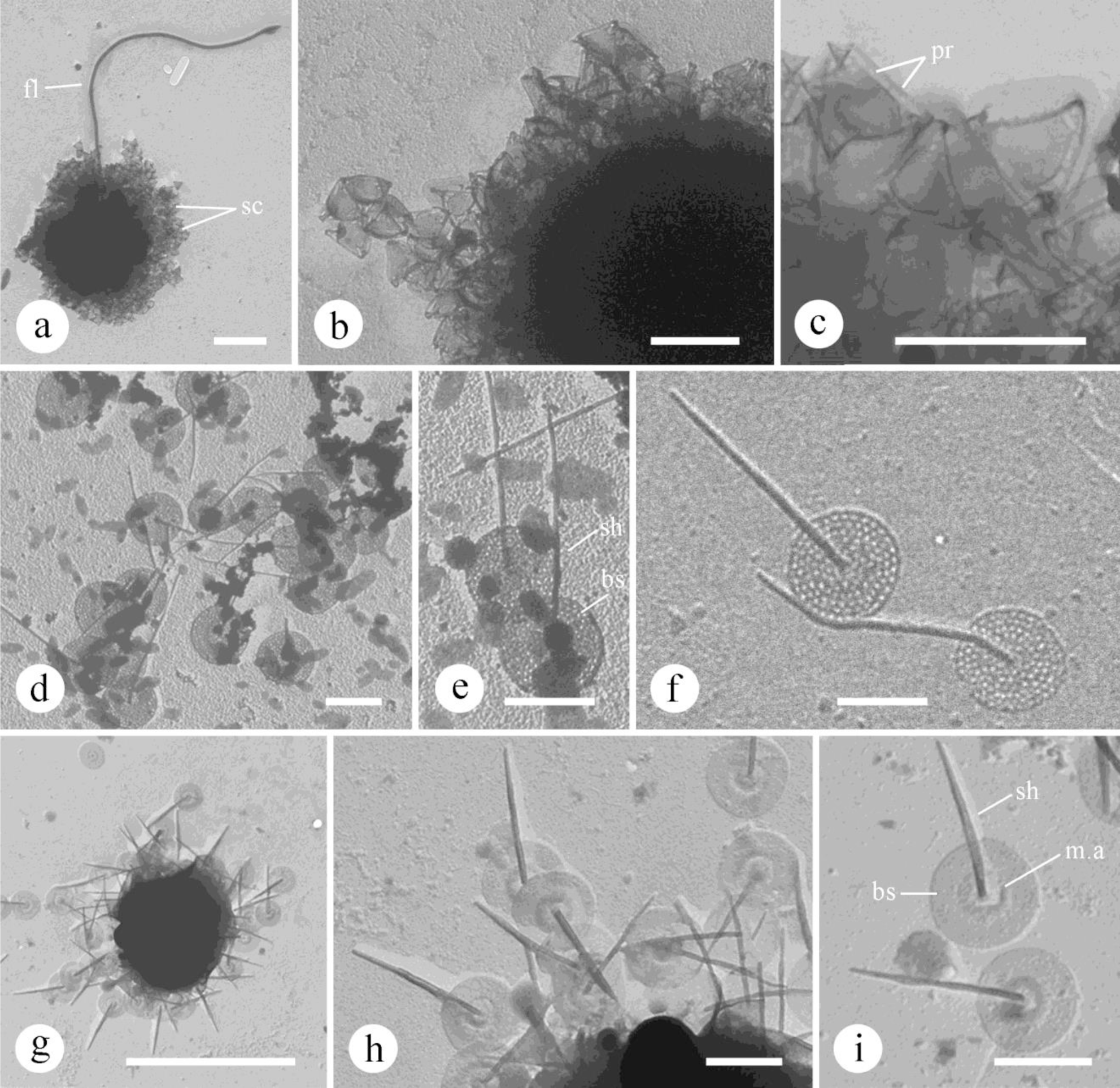
Morphology of observed heterotrophic flagellates (TEM): a–c – *Thaumatomonas seravini*; d–f – *Paraphysomonas foraminifera*; g–i – *Paraphysomonas* sp. Abbreviations: bs – base of spine scale; fl – flagellum; m.a – middle annulus; sc – scales; sh – shaft; pr – perforations. Scale bars: a, g – 5 µm; b, c, d–f, h, I – 1 µm.

### Observations

Living cells were not observed. According to TEM whole mount, protoplast is ovate, 6.5–7.5 μm long and 4.5–6.0 μm wide. Long flagellum is 20–23 μm in length, short flagellum wasn’t observed. Cells covered by numerous siliceous scales of one type. Scales consist of two triangular plates, connected to each other with three hollow cylindrical struts at the corners (0.2–0.3 μm in length). Sides of plates are about 0.5–0.6 μm in length. Inner plate textureless, with more rounded corners. External plate with gently curved and more pointed corners and row of perforations along margin. A large oval hole located in each corner. Observations based on description of scales of five cells in TEM.

### Remarks

This species is common in freshwater, but is extremely rare in marine waters (see distribution). Our observations are corresponded with previous findings, except authors describe larger sizes of the cell: length is 8.4–15.0 μm, width is 5.0–13 μm (Mylnikov and Karpov 1993; Mylnikov et al. 2002, 2006; Prokina et al. 2017; Tikhonenkov 2006). Cell protoplasts could shrink when dried. Size and morphology of observed scales are similar to described previously. This species is quite similar to *T. hindoni* (Nicholls, 2013) Scoble et Cavalier-Smith, 2014, described from fresh waters of Canada (Nicholls 2013). *T. seravini* differs from *T. hindoni* by larger perforations along plate margins, presence of a large oval hole at each corner of external plate, and more gentle curved corners (Mylnikov and Karpov 1993; Nicholls 2013; Scoble and Cavalier-Smith 2014b). Another species of the genus *Thaumatomonas* with similar scales is *T. zhukovi* Mylnikov, 2003, can be distinguished by the presence of two types of scales – oval and triangular.

However, Scoble and Cavalier-Smith (2014b) studied the morphology and 18S rRNA sequences of several clones of this species and found that this species can be polymorphic with two or one type of scales (either oval or triangular), so these species cannot be distinguished by the morphology of scales with 100% certainty. Their studies also have shown that both species are closely related.

### Distribution

Fresh waters of European part of Russia (Kopylov et al. 2015;Mylnikov and Karpov 1993; Mylnikov et al. 2002, 2006; Prokina et al. 2017; Tikhonenkov 2006), Mongolia (Kopylov et al. 2006), Chile (Prokina and Mylnikov 2018), China (Tikhonenkov et al. 2012). Soils of China (Tikhonenkov et al. 2012). White Sea (Tikhonenkov 2006; Tikhonenkov and Mazei 2013).

***Metromonadea Cavalier-Smith, 2007

*Metopion fluens* Larsen et Patterson, 1990^c^ – observed in sample points 2a, 2b, 3c, 4b, 5b, 5c, 6, 7, 8a.

*Metromonas grandis* Larsen et Patterson, 1990^c^ – observed in sample points 2a, 2b, 5b, 6.

M. simplex (Griessmann, 1913) Larsen et Patterson, 1990c – observed in sample points 2a, 2b, 7.

***Granofilosea Cavalier-Smith et Bass, 2009

****Massisteridae Cavalier-Smith, 1993

*Massisteria marina* Larsen et Pattersen, 1990^c^ – observed in sample points 2a, 4d, 5b.

***Glissomonadida Howe et Cavalier-Smith, 2009

**** Dujardinidae Howe et Cavalier-Smith, 2011

****Allapsidae Howe et Cavalier-Smith, 2009

*Allantion tachyploon* Sandon, 1924^c^ – observed in sample point 3a.

*Incertae sedis* Cercozoa

*Discocelis punctata* Larsen et Patterson, 1990^c^ – observed in sample points 5b, 7.

*Stramenopiles Patterson, 1989

**Bigyra Cavalier-Smith, 1998

***Opalozoa Cavalier-Smith, 1991

****Bicosoecida Grasse, 1926

*Bicosoeca gracilipes* James-Clark, 1867^c^ – observed in sample point 2b.

*B. maris* Picken, 1841^c^ – observed in sample points 1b, 8b.

*Caecitellus parvulus* (Griessmann, 1913) Patterson et al., 1998^c^ – observed in sample points 2a, 2e, 4d, 5c, 6, 7, 8b.

*Cafeteria ligulifera* Larsen et Patterson, 1990^c^ – observed in sample points 3a, 4a, 5b, 6. *C. minuta* (Ruinen, 1938) Larsen et Patterson, 1990^c^ – observed in sample points 4d, 5c. *C. roenbergensis* Fenchel et Patterson, 1988^c^ – observed in sample points 2a, 2b, 2c, 3a, 3b, 4a, 4b, 5a, 5b, 5c, 6, 7.

*Halocafeteria seosinensis* Park et al., 2006 – observed in sample point 8b.

*Pseudobodo tremulans* Griessmann, 1913^c^ – observed in sample point 6.

**Gyrista Cavalier-Smith, 1998

***Ochrophyta Cavalier-Smith, 1986

****Diatomista Derelle et al., 2016

*****Dictyochophyceae Silva, 1980

******Pedinellales Zimmermann et al., 1984

*Actinomonas mirabilis* Kent, 1880–1882^c^ – observed in sample point 4d. *Ciliophrys infusionum* Cienkowski, 1876^c^ – observed in sample points 5c, 6, 8b. *Pteridomonas danica* Patterson et Fenchel, 1985^c^ – observed in sample points 3b, 3c, 4d, 5a.

****Chrysista Cavalier-Smith, 1986

*****Chrysophyceae Pascher, 1914

******Paraphysomonadida Scoble et Cavalier-Smith, 2014

*Clathromonas butcheri* (Pennick et Clarke, 1972) Scoble et Cavalier-Smith, 2014^c^ – observed in sample point 3a.

***Paraphysomonas foraminifera*** Lucas, 1967^c^ (Fig. 3d–f) – observed in sample points 3b, 6. <u>Observations</u>

Living cells were not observed. Cells covered by siliceous hobnail-like scales of one type. Scales consist of circular flattened base and cylindrical shaft. Base is 0.7–1.2 μm in diameter, perforated with 10 or more concentric rows of round holes, which are not always clearly visible. Shaft is 1.9–3.0 μm in length, 0.05–0.1 μm in diameter, with sharp end. Several single scales and scale aggregations of two cells observed in TEM.

### Remarks

Many authors described smaller length of shaft, 0.45–1.3 μm (Bergesch et al. 2008; Lukas 1967; Tong 1997b, c; Vørs 1993); 7–8 concentric rows of holes on base (LeRoi and Hallegraeff 2006; Tong 1997c;Tong et al. 1998), as well as presence of middle dense annulus (LeRoi and Hallegraeff 2006; Tong 1997c), and “sholders” – sharp narrowing of diameter of the shaft in the middle of its length (Tong 1997c). This species differs from others by presence of 7–10 concentric rows of holes on base plate and shaft without perforations on its basal part. Nicholls (1981) described several scales from freshwater in Canada, under the name *P. foraminifera*.

However, these scales probably belong to several *Paraphysomonas* or *Clathromonas* species because of significant differences: presence of perforations on the basal part of shaft, as well as reduced shaft on some scales, larger size of scales (base up to 2.5 μm). Several scales (Nicholls 1981: Fig. 16–18, p. 134) are very similar to scales of species *Clathromonas takahashii* (Cronberg et Kristiansen, 1981) Scoble et Cavalier-Smith, 2014 and one scales (Nicholls 1981: Fig. 19, p. 134) are very similar to scales of *Clathromonas elegantissima* (Kling et Kristiansen, 1983) Scoble et Cavalier-Smith, 2014.

### Distribution

Marine waters of U.K. (Tong 1997b), Croatia (Leadbeater 1974), Brazil (Bergesch et al. 2008), Canada (Smith and Hobson 1994), Australia (LeRoi and Hallegraeff 2006; Tong 1997c; Tong et al. 1998), Greenland (Vørs 1993), Baltic Sea (Vørs 1992), MediterraneanSea (Throndsen and Zingone 1994).

***Paraphysomonas* sp.** (Fig. 3g–i) – observed in sample point 3d.

### Observations

Living cells were not observed. Cells are covered by siliceous hobnail-like scales of one type. Scales consist of circular flattened base and cylindrical shaft. Base is 0.6–0.9 μm in diameter, with a dense middle annulus. Shaft is 1.3–1.9 μm in length, 0.08–0.1 μm in diameter, conically pointed into a rounded tip. Scales of two cells observed in TEM.

### Remarks

Scales of this species are most similar to scales of *P. imperforata* Lukas, 1967 by small length of shaft, base plate with middle annulus, and lacking dense margins. But observed scales are differentiated by a noticeably thinner shaft without conical tapering. Other species with middle annulus on base plate (*P. acuminata acuminata* Scoble et Cavalier-Smith, 2014; *P. acuminata cuspidata* Scoble et Cavalier-Smith, 2014; *P.mikadiforma* Scoble et Cavalier-Smith, 2014; *P*. *hebetispina hebetispina* Scoble et Cavalier-Smith, 2014) differ due to a very long shaft relative to base plate diameter (Scoble and Cavalier-Smith 2014a).

Cryptista Adl et al., 2018

*Cryptophyceae Pascher, 1913

**Cyathomonadacea Pringsheim, 1944

*Goniomonas pacifica* Larsen et Patterson, 1990^c^ – observed in sample points 2a, 3d, 6, 7.

G. truncata (Fresenius, 1858) Stein, 1878c – observed in sample points 2a, 2d, 3b, 4d, 5c, 6, 7.

*Goniomonas* sp. – observed in sample point 8a.

Haptista Cavalier-Smith, 2003

*Centroplasthelida Febvre-Chevalier et Febvre, 1984

**Panacanthocystida Shɨshkin et Zlatogursky, 2018

****Acanthocystida Cavalier-Smith et von der Heyden, 2007

*****Chalarothoracina Hertwig et Lesser, 1874 *sensu* Cavalier-Smith, 2012

******Raphidocystidae Zlatogursky, 2018

***Raphidocystis bruni*** (Penard, 1903) Zlatogursky, 2018 (Fig. 4a–c) – observed in sample point 2a.

**Fig. 4.**
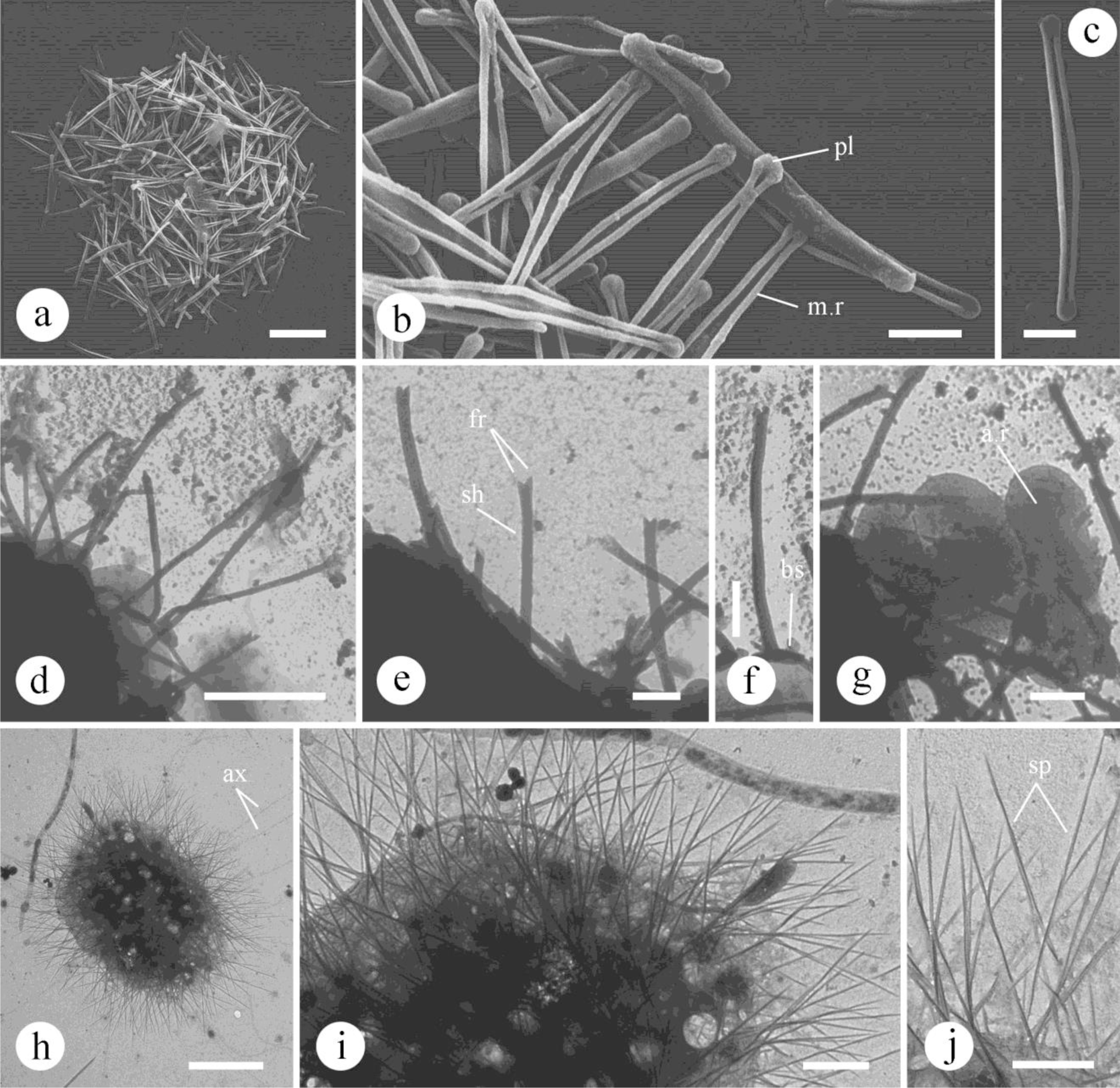
Morphology of observed scales of centrohelid heliozoans (a–c – SEM; d–j – TEM): a–c – *Raphydocystis bruni* (a, b – scale aggregation of whole cell; c – single scale); d–g – *Choanocystis perpusilla* (d, e – scale aggregation of whole cell; f – spine scale; g – plate scales); h–j – *Heterophrys-*like organism. Abbreviations: a.r – axial ridge; ax – axopodia; bs – base of scale; fr – furcae; m.r – marginal rim; pl – poles of scales; sh – shaft; sp – spicules. Scale bars: a, d, h – 5 µm; b, c, e–g, I, j – 1 µm.

### Synonyms

*Raphidiophrys bruni* Penard, 1903; *Polyplacocystis brunii* (Penard, 1903) Mikrjukov, 1999.

### Observations

Cells covered by tangentially-oriented elongated-oval plate scales of one type. Scales are 5.9–8.1 μm in length, that are slightly widened in the central part (0.5–0.8 μm) and gradually taper to the poles (where they are 0.3–0.4 μm), forming circular poles 0.5–0.6 μm in width. The length to width ratio is 8.2–13.8. Inner surface of scales is smooth, without a longitudinal rib. Hollow marginal rim is 0.1–0.2 μm in diameter in the middle, and expands to the poles. Scales of one cell were observed in SEM.

### Remarks

This species can be distinguished from other species of *Raphidocystis* due to elongated shape and circular poles, the lack of reticular structure on the inner surface of scales, and the absence of spine scales.

### Distribution

Fresh waters of Crimea (Mikrjukov 1999). Black Sea (Mikrjukov 1999), Tasman Sea (Mikrjukov 2002).

**Pterocystida Cavalier-Smith et von der Heyden, 2007

***Raphidista Shɨshkin et Zlatogursky, 2018

****Choanocystidae Cavalier-Smith et von der Heyden, 2007

***Choanocystis perpusilla*** (Petersen et Hansen, 1960) Siemensma, 1991^c^ (Fig. 4d–g) – observed in sample point 8b.

### Synonyms

Acanthocystis perpusilla Petersen et Hansen, 1960; Choanocystis cordiformis ssp*. parvula* Dürrschmidt, 1987.

### Observations

Cells covered by spine and plate scales. Radial spine scales consist of hollow cylindrical shaft, asymmetrically extending from the heart-shaped flattened base. Shaft is slightly curved, 2.7–8.0 μm in length, 0.15–0.2 μm in diameter, and the distal tip divides on two furcae. Base is 1.1–1.7 μm in diameter, with a small notch. Tangential oval plate scales are 1.5–2.5×1.5– 2.0 μm, with concave long sides and an axial ridge located inside medial depression. We observed scales of four cells in TEM.

### Remarks

The size of observed scales corresponds with scales of subspecies *Ch*. *cordiformis parvula*, described by Dürrschmidt (1987a). Most often a shorter length of spine scales is indicated in the literature, 1.4–5.6 μm (Croome et al. 1987; Dürrschmidt 1985; Shatilovich et al. 2010), however, there are records with a length 8–10 μm (Leonov and Mylnikov

2012). Also, we previously described textureless plate scales, without axial ridge (Prokina and Mylnikov 2019).

### Distribution

Freshwater of European part of Russia (Leonov and Mylnikov 2012), Estonia (Mikrjukov 1993), Chile (Dürrschmidt 1985; Prokina and Mylnikov 2019), Sri Lanka ([as *Acanthocystis cordiformis parvula*] Dürrschmidt 1987), Vietnam (Prokina et al. 2019). Soil of East Siberia, Russia (Shatilovich et al. 2010). Marine waters of Antarctica (Croome et al. 1987). Baltic Sea (Vørs 1992). Saline inland waters of European part of Russia (Ermolenko and Plotnikov 2013).

***Pterista Shɨshkin et Zlatogursky, 2018

*\*\*\*\**Heterophryidae Poche, 1913

***Heterophrys***-like organism (Fig. 4h–j) – observed in sample point 5c.

### _Observations_

Cells covered by radial organic fusiform spicules 5.2–11.5 μm in length, 0.05–0.1 μm in wide. Spicules are slightly flattened and spirally twisted along the longitudinal axis, and both tips are pointed. Basal part of spicules immersed in mucopolysaccharide capsule that is sometimes visible in TEM, as are axopodia. Spicules of five cells were observed in TEM.

### Remarks

Centrohelids with spindle-shaped organic spicules have previously been found from various freshwater and marine habitats around the world, and have been identified as species of genera *Heterophrys*, *Marophrys*,and *Sphaerastrum* (Cavalier-Smith and von der Heyden 2007; Mikrjukov 2002). However, recent studies have revealed close relations between spicule-bearing *Heterophrys*-like organisms and siliceous scales-bearing species both by molecular data (18S rDNA gene similarity of *Polyplacocystis* and *Heterophrys*), and by morphological data: intermediate polymorphic stages of the life cycle of *Raphidiophrys*, bearing both siliceous scales and organic spicules (Zlatogursky 2016; Zlatogursky et al. 2018). Based on these observations, Zlatogursky with coauthors (Zlatogursky et al. 2018) suggested dimorphism of centrohelid life cycle with two stages, one with siliceous scales and another with organic spicules. Thus, spicules of the cells we studied may be the stage of the life cycle of one or more scale-bearing centrohelid species.

*Incertae sedis* Diaphoretickes

Telonemia Shalchian-Tabrizi, 2006

*Telonema subtile* Griessmann, 1913^c^ – observed in sample point 2b, 6.

Incertae sedis EUKARYA: EXCAVATES [Excavata Cavalier-Smith, 2002, emend. Simpson, 2003]

*Discoba Simpson, 2009

**Euglenozoa Cavalier-Smith, 1981

***Kinetoplastea Honigberg, 1963

*Incertae sedis* Kinetoplastea

*Pseudophyllomitus apiculatus* (Skuja, 1948) Lee, 2002 *sensu* Mylnikov, 1986^c^ – observed in sample points 2a, 2b, 2d, 3a, 3b, 5b.

****Metakinetoplastina Vickerman, 2004

*****Neobodonida Vickerman, 2004

*Neobodo designis* (Skuja, 1948) Moreira et al., 2004^c^ – observed in sample points 2a, 2c, 2d, 3a, 3b, 4a, 4b, 5b, 6, 7, 8a.

*N. saliens* (Larsen et Patterson, 1990) Moreira et al., 2004^c^ – observed in sample points 2a, 3a, 4b, 5b, 6, 8a.

*Rhynchomonas nasuta* (Stokes, 1888) Klebs, 1893^c^ – observed in sample points 2a, 2c, 2d, 3a, 3b, 3c, 3d, 4a, 4b, 4c, 4d, 5b, 5c, 6, 7, 8a, 8b.

*Rhynchobodo simius* Patterson et Simpson, 1996^c^ – observed in sample point 6.

***Euglenida Butschli, 1884

****Heteronematina Leedale, 1967

*Dinema platysomum* (Skuja, 1939) Lee et Patterson, 2000^c^ – observed in sample points 6.

*Lentomonas azurina* (Patterson et Simpson, 1996) Cavalier-Smith, 2016^c^ – observed in sample points 2a, 2b, 5b, 5c, 6, 7.

*L. corrugata* (Larsen et Patterson, 1990) Cavalier-Smith, 2016^c^ – observed in sample points 2a, 6, 7.

***Petalomonas cantuscygni*** Cann et Pennick, 1986 (Fig. 5a–c) – observed in sample points

1a, 1b, 3a.

**Fig. 5.**
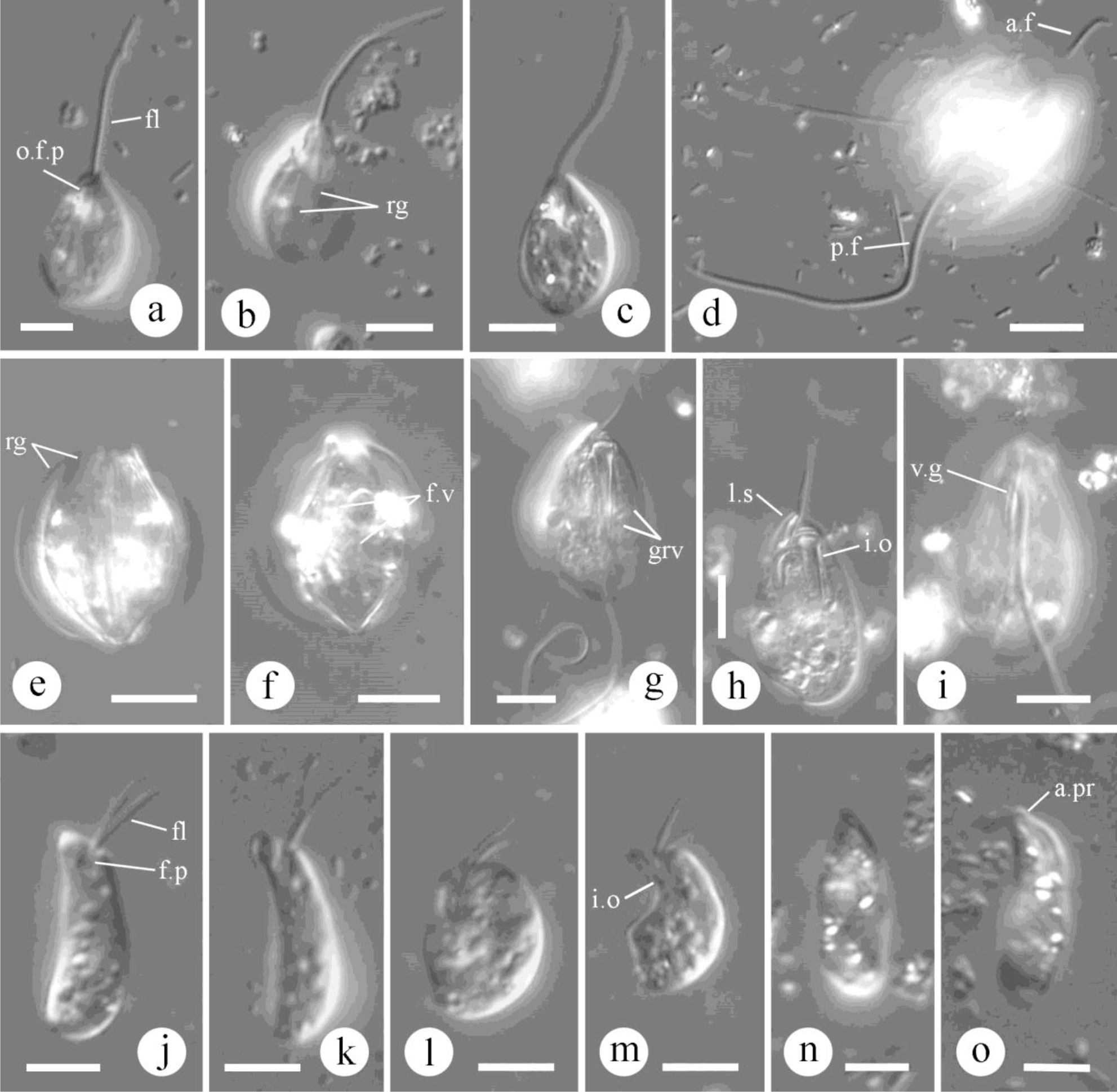
Morphology of observed heterotrophic flagellates (DIC): a–c – *Petalomonas cantuscygni*; d–f – *Ploeotia adhaerens*; g–i – *P. robusta*; i–m – *Diplonema ambulator*; n, o – *Rhynchopus amitus*. Abbreviations: a.f – anterior flagellum; a.pr – apical protrusion (papilla); a.sl – anterior sleeve; fl – flagellum; f.p – flagellar pocket; f.v – food vacuole; grv – grooves; i.o – ingestion organelle; l.s – lip-like structure; p.f – posterior flagellum; p.sl – posterior sleeve; v.g – ventral groove; o.f.p – openings of flagellar pocket; rg – ridges. Scale bars: a–c, j–o – 5 µm; d–i – 10 µm.

### Observations

Cells are ovoid in outline, anterior end slightly narrowed with well-marked openings of flagellar pocket (Fig. 5a). Cell length is 9.5–14.0 μm, width is 6–9 μm. Single flagellum is about the cell length or slightly longer, directed anteriorly. Dorsal side of the cell with 6 gentle ridges (Fig. 5b), ventral side with two slightly noticeable ridges. Flagellar pocket at the right side of the cell. Nucleus located in the anterior part of the cell. Observations based on four cells found in LM.

### Remarks

Our observations completely correspond with previous descriptions of this species. Cell length is 16 μm (Larsen and Patterson 1990) and 9.2–15.5 μm in the original description (Cann and Pennick 1986). The other species of the genus *Petalomonas* all have oval cell outlines (*P*. *poosilla*, *P. ingosus* Lee et Patterson, 2000, *P*. *labrum* Lee et Patterson, 2000, *P*. *minor*, *P*. *minuta*, *P*. *tricarinata* Skuja, 1939, *P. vigatus* Lee et Patterson, 2000), and differ in number or noticeability of dorsal and ventral ridges, or by the presence of a smooth cell surface.

### Distribution

Marine waters of U.K. (Cann and Pennick 1986), Brazil (Larsen and Patterson 1990).

1. *P. minor* Larsen et Patterson, 1990^c^ – observed in sample points 1b, 2a, 3a, 6, 7.
2. *P. minuta* Hollande, 1942^c^ – observed in sample points 5a, 5b, 6, 7.
3. *P. ornata* Skvortzov, 1957^c^ – observed in sample point 3d.
4. *P. poosilla* (Skuja, 1948) Larsen et Patterson, 1990^c^ – observed in sample points 2a, 3a, 3b, 4d, 5a, 5b, 6, 7, 8b.

*Petalomonas* sp. – observed in sample point 3c.

***Ploeotia adhaerens*** Larsen et Patterson, 1990^c^ (Fig. 5d–f) – observed in sample points 6, 7. <u>Observations</u>: Cells not flattened, almost circular in outline, with four prominent dorsal ridges, two lateral ridges, and two ventral ridges. Posterior end of the cell is slightly extended into a small protrusion. Cell length is 26–30 μm, width is 22–27 μm. Anterior flagellum in about half of the cell length. Posterior flagellum is about 2.5 time longer than the cell, and noticeably thicker than the anterior flagellum. Flagellar pocket is situated on the left side of the cell and the nucleus on the right side. Cells contains large food vacuoles in their posterior half. Well-marked ingestion organelle with two rods located on the right side, and broad anteriorly. Observed seven cells in LM.

### Remarks

Morphology of observed cells corresponds with previous descriptions, except longer anterior flagellum (equal to the cell length) and shorter posterior flagellum (about 1.5 times longer than the cell length) (Ekebom et al. 1995/6; Larsen and Patterson 1990). This species is easily distinguished from other *Ploeotia* species by almost round cells in outline and very prominent ridges.

### Distribution

Marine waters of Australia (Ekebom et al. 1995/6), Fiji (Larsen and Patterson 1990), Red Sea (Tikhonenkov 2009). Fresh waters of European part of Russia (Tikhonenkov 2006; Tikhonenkov and Mazei 2007).

1. *P.* aff. *costata* (Triemer, 1986) Farmer et Triemer, 1988^c^ – observed in sample point 7.
2. *P. discoides* Larsen et Patterson, 1990^c^ – observed in sample points 5b, 7. *P. oblonga* Larsen et Patterson, 1990^c^ – observed in sample points 2a, 8a. *P. punctata* Larsen et Patterson, 1990 – observed in sample point 3a.

***Ploeotia robusta*** Larsen et Patterson, 1990 (Fig. 5g–i) – observed in sample point 3b. <u>Observations</u>: Cells oval in outline, with subapically pointed posterior end (Fig. 5h). Cell length is 30–35 μm, cell width is 20–22 μm. Flagellar pocket opened subapically and covered laterally by lip-like flange (Fig. 5h). Anterior flagellum is about the cell length. Posterior flagellum is 3.5 times longer than the cell, and wider than anterior flagellum. One well-developed ventral groove extends from lip-like structure to the posterior pointed tip (Fig. 5i), and about 4–5 slightly noticeable grooves situated on both ventral and dorsal sides of the cell (Fig. 5g). Well-marked ingestion organelle with two gradually narrowing rods extends along entire cell length. Nucleus located at the left side of the cell. Posterior part of the cell contains a few food vacuoles. Cells lack any surface structure, except the ventral groove associated with lip-like flanges, and a rough surface. Four cells were found in LM.

### Remarks

We observed about 8–10 poorly visible grooves on cell body surface, despite these not being part of the original description by Larsen and Patterson (1990). Patterson and Simpson (1996) also noticed at least eight delicate grooves arranged evenly around the cell body, which they described as being faintly visible and easily overlooked. At the same time, Al-Qassab with coauthors (2002) recorded 3–4 ventral ridges and 4–5 dorsal ridges. Patterson and Simpson 1. (1996) described longer flagella, with the anterior flagellum twice as long as the cell and posterior flagellum 4 times the length of the cell. This species differs from other *Ploeotia* species by larger cell size, presence of anterior-lateral lip-like flanges with extended well-developed ventral groove, long posterior flagellum, and posterior pointed end of the cell.

### Distribution

Marine waters of U.K., Hawaii (Larsen and Patterson 1990). Hypersaline water of Australia (Al-Qassab et al. 2002; Patterson and Simpson 1996).

*Ploeotia* sp. 1 – observed in sample points 2a, 5a, 5b, 5c.

*Ploeotia* sp. 2 – observed in sample points 2a, 2b, 3c, 5a, 5b, 6, 8a.

***Diplonemea Cavalier-Smith, 1993

****Diplonemidae Cavalier-Smith, 1993

***Diplonema ambulator*** Larsen et Patterson, 1990^c^ (Fig. 5j–m) – observed in sample points 2c, 5b.

### Observations

Cells are highly metabolic, can twist and contract to almost rounded in outline during turns (Fig. 5l). Cells are 14.5–16 μm long, 4–6 μm wide, elliptical with slightly rounded posterior end and obliquely cut anterior end. Equal-sized thin flagella emerge subapically from prominent flagellar pocket, length of flagella is 5–6 μm. Flagella usually direct posteriorly or slowly alternately flex to the lateral side giving the impression of “walking legs”. J-shaped ingestion organelle seen during metaboly (Fig. 5l, m). Cell surface is bumpy. Only gliding cells were observed. Observations based on findings of six cells in LM.

### Remarks

Morphology of studied cells mostly corresponds with previous descriptions, except that shorter flagella were described by Larsen and Patterson (1990), 3–4 μm; and longer flagella were described by Tong (1994, 1997b), 8 μm. Also, Tong (1994) noted that one flagellum sometimes can wraps around the cell, but we did not observe this. Other known species of this genus, *D. breviciliata* Griessmann, 1914, and *D. metabolicum* Larsen et Patterson, 1990, can be distinguished by cells being twice as large (Tong et al. 1998); *D. aggregatum* Tashyreva et al., 2018 and *D. japonicum* Tashyreva et al., 2018 can be distinguished by broader anterior part of the cell and narrowed anterior end, as well as presence of a sessile rounded cell stage and swimming stage with unequal flagella twice the length of the cell (Tashyreva et al. 2018); *D. papillatum* (Porter, 1973) Triemer et Ott, 1990 can be distinguished by oval to almost round outline of cells, the presence of prominent apical papilla, and very short, thick flagella (Porter 1973). Morphology of studied species is similar to *Rhynchopus amitus* by its cell size and shape, but latter species can be distinguished by presence of flagella that are thicker, barely emerge from flagellar pocket, curve posteriorly, and are mostly motionless (Al-Qassab et al. 2002; Tong et al. 1998), see below.

### Distribution

Marine waters of Australia (Tong et al. 1998), Brazil (Larsen and Patterson 1990), U.K. (Patterson et al. 1993; Tong 1994, 1997c).

***Rhynchopus amitus*** Skuja, 1948^c^ (Fig. 5n, o) – observed in sample points 2a, 8b. <u>Synonym</u>: *R. conscinodiscivorus* Schneps, 1994.

### Observations

Cells sac-shaped, with rounded ends and fine anterior apical papilla. Cell length is 9.5–12 μm, width is 4–5.5 μm. Cells very metabolic, periodically contracts. Flagellar pocket opened below the apical papilla, directed anterior-laterally. Flagella usually do not emerge from pocket. Numerous granules are visible inside the cell just below the cell surface. Cells glide slowly, constantly change directions; swimming cells were not observed. Two cells were found in LM.

### Remarks

Most authors observed larger cells, up to 25 μm (Al-Qassab et al. 2002; Lee, 2015; Schnepf 1994; Skuja 1948; Tong et al. 1998). Studied species differs from other species of *Rhynchopus* by cell shape and size. *R. euleeides* Roy et al., 2007 differs by presence of long unequal flagella in swimming cells (2.5 times longer than the cell), symmetric elliptical shape of resting cells, clusters of many cells in culture (Roy et al. 2007). *R. hemris* Tashyreva et al., 2018 differs by both narrowed ends of the cell, swimming cells with one flagellum wrapped around anterior part of the cell and wobbling like a lasso, and another flagellum extended along the cell and waving (Tashyreva et al. 2018). *R. serpens* Tashyreva et al., 2018 has tapered anterior part of the cell and broad posterior end, as well as larger cell size (Tashyreva et al. 2018). Observed species is also similar to *Diplonema ambulator* (see above).

### Distribution

Marine waters of Australia (Lee 2015; Lee et al. 2003; Tong et al. 1998), Wadden Sea ([as *R. conscinodiscivorus*] Schneps 1994). Saline inland waters of Romania ([as *Menoidium astasia*] Entz 1883). Hypersaline waters of Australia (Al-Qassab et al. 2002; [as *Menoidium astasia*] Ruinen 1938). Fresh waters of Sweden (Skuja 1948).

**Heterolobosea Page et Blanton, 1985

***Tetramitia Cavalier-Smith, 1993

****Eutetramitia Hanousková et al., 2018

*****Percolomonadidae Cavalier-Smith, 2008

*Percolomonas cosmopolitus* (Ruinen, 1938) Fenchel et Patterson, 1986^c^ – observed in sample points 1a, 7.

1. *P. denhami* Tong, 1997^c^ – observed in sample points 1b, 2a, 2c, 5b, 6.
2. *P. similis* Lee et al., 2003^c^ – observed in sample points 1a, 2a.

*Incertae sedis* Eukarya: CRuMs

*“CRuMs” (Brown et al., 2018)

*Glissandra innuerende* Patterson et Simpson, 1996^c^ – observed in sample point 6.

*Incertae sedis* Eukarya: CRuMs *CRuMs” (Brown et al. 2018) [Varisulca Cavalier-Smith 2012] (R) *\*\*Mantamonas* Cavalier-Smith et Glücksman, 2011

***Mantamonas plastica*** Glücksman et Cavalier-Smith in Glücksman et al., 2011 (Fig. 6a–c)

**Fig. 6.**
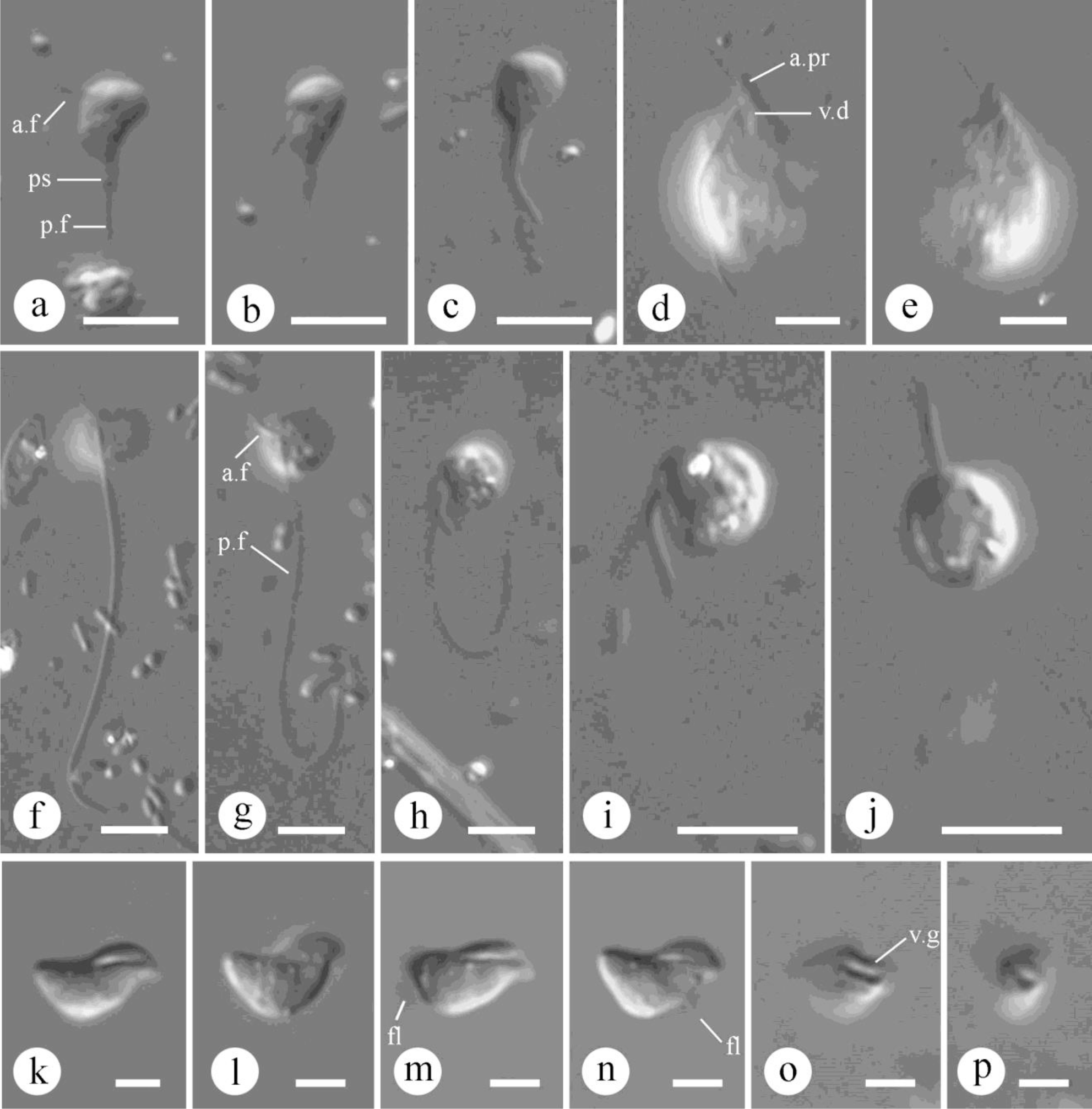
Morphology of observed heterotrophic flagellates (DIC): a–c – *Mantamonas plastica*; d, e – “*Heterochromonas opaca*”; f–i – “Protist 1”; (f – both flagella in focus; g, h – cell turns; i, j – cell and anterior flagellum in focus); k–p – “Protist 2” (k–n – view from lateral side; o, p – view from ventral side, with ventral groove). Abbreviations: a.f – anterior flagellum; a.pr – apical protrusion; fl – flagellum; p.f – posterior flagellum; ps – pseudopodia; v.d – ventral depression; v.g – ventral groove. Scale bars: 5 µm.

– observed in sample points 3a, 6, 7.

### Observations

Cells ovoid or triangular, highly metabolic, with posterior end extended into a changeable tail pseudopodium, connected with posterior flagellum. Left side of the cell is smaller than right side. Cell length is 3.5–4.5 μm, width is 2.5–3.0 μm at the broadest part.

Anterior flagellum is thin, directed laterally (to the left), and moves rarely. Length of anterior flagellum is equal to the cell length. Posterior flagellum is about twice the length of the cell, thicker than the anterior flagellum, and trails behind the cell. Both flagella non-acronematic. Ten cells were observed in LM.

### Remarks

Size and morphology of observed cells are similar to those observed earlier, except in original description authors noted acronematic posterior flagellum (Glücksman et al. 2011). This species differs from other flagellates by its small size of cells, lateral orientation of anterior flagellum and flexible triangular shape of cells.

### Distribution

Marine waters of U.K. (Glücksman et al. 2011), Australia (Lee 2015).

*Incertae sedis* Eukarya: Ancyromonadida Cavalier-Smith, 1998

*Ancyromonas sigmoides* (Kent, 1880–1882) Heiss et al., 2010^c^ – observed in sample points 2a, 3a, 3d, 5b, 6, 7, 8b.

Other *incertae sedis* Eukarya

**“*Heterochromonas opaca*”** *sensu* Lee et Patterson, 2000 (Fig. 6d, e) – observed in sample point 6.

### Observations

Oval cells with slightly narrowed ends and apical anterior protrusion. Cell length is 11–15 μm, cell width is 8.5–11.0 μm. Flagella emerge from the ventral subapical depression at the base of apical protrusion. Anterior flagellum is half the length of the cell, directed anterior-laterally, and mostly does not move. Posterior flagellum is 1.1–1.2 times the length of the cell, located laterally in a curve, and trailed behind the cell. Nucleus located anteriorly. Surface of cells is warty. Cells move in a wide circle in a clockwise direction. Three cells were observed in LM.

### Remarks

Morphology of cells we observed agreed well with cells described by Lee and Patterson (2000) as “*Heterochromonas opaca*”. However, despite the similar cell shape, the cell originally described as *Heterochromonas opaca* Skuja (1948) is probably a different organism. The genus *Heterochromonas* Pascher, 1912 was introduced for colourless chrysomonads without scales, related to *Ochromonas* Vysotskii, 1887, and, probably, a junior synonym of *Spumella* Cienkowski, 1870 (Grossmann et al. 2016). Species of *Heterochromonas* are characteried by a clearly stramenopile body plan, with non-flattened metabolic cells, truncated apically, and with two unequal heterodynamic flagella, both directed anteriorly, and with small protoplasmic collar around apical depression. Species *Heterochromonas opaca*, described by Skuja (1948) are characterized by globular cells with well-developed amoeboid collar that can almost completely disappear (except for the basal part), and the presence of short finger-like irregular pseudopodia involved in the capture of food particles. The cell described by Lee and Patterson (2000) as

“*Heterochromonas opaca*” cannot be attributed to Chrysophyceae due to rigid and flattened cells without an anterior collar and pseudopodia, or the long flagellum directed posteriorly and trailed behind the cell. In addition, Skuja described freshwater species with 2–3 contractile vacuoles, while “*Heterochromonas opaca*” *sensu* Lee et Patterson, 2000 was found only in marine waters (see distribution). Thus, most likely Lee and Patterson described a completely different species, probably new, that needs to be renamed and investigated to clarify its relationships with other eukaryotes. According to the external morphology and movement, this organism could tentatively be attributed to Euglenida.

Previously described cells from the Black Sea (Prokina et al. 2018), identified as *Heterochromonas opaca* Skuja, 1948, are characterized by markedly different morphology: circular cells in outline; poorly developed anterior protrusion; smaller cell sizes; greater length and mobility of flagella. Probably it was another, undescribed species, related to “*Heterochromonas opaca*” *sensu* Lee et Patterson, 2000.

### Distribution

Marine waters of Australia (Lee and Patterson 2000), South Korea (Lee 2002).

**“Protist 1”** (Fig. 6f–j) – observed in sample point 7.

### Observations

Rigid cells are either spherical or ovoid, with slightly flattened ventral side.

Cell length is 4.5–5.5 μm, cell width is 3.8–5.5 μm. Unequal non-acronematic flagella insert subapically from ventral side. Anterior flagellum is about the cell length, directed mostly forward, without moving, but can turn back when the cell turns (Fig. 6g, h). Posterior flagellum is about 25–35 μm long, trailed behind the cell. Pseudopodia were not observed. Nucleus located medially.

Cells move slowly, gliding on the posterior flagellum. Four cells were observed in LM.

### Remarks

Observed organism differed from other known flagellate species by presence of very long posterior flagellum relative to the cell length. Long posterior flagellum and gliding cells are typical for *Glissandra innuerende* and *G. similis* Lee, 2006. But listed species are differed by the following morphological features: longer anterior flagellum which almost equal in length to posterior flagellum; flagella insert from subapical depression; cells glide on both flagella, and anterior flagellum always directed anteriorly; *G. similis* has ventral longitudinal groove (Al-Qassab et al. 2002; Lee 2006). Observed unidentified protist is also similar to *Sinistermonas sinistrorsus* Lee, 2015 which is distinguished from our taxon by its longer flagellum, directed anterior-laterally and beating always to left in a small excursion (Lee 2015).

**“Protist 2”** (Fig. 6k–p) – observed in sample point 7.

### Observations

Cells rigid, triangular in outline, laterally flattened, with tapered and curved ends, dorsally convex and ventrally concave with large longitudinal groove that widens to one end of the cell. Cell length is 4.5–6.0 μm, width is 3.0–3.5 μm. Cells also slightly curved in longitudinal axis (Fig. 6o). Flagella usually not seen, except possibly through perturbations of water at the edges of the cell. Sometimes a very thin flagellum (or filopodium?) is visible under the cell (Fig. 6m, n), which makes circular movements, potentially helping the cell move. Cells are located near a substrate and rotate around its axis. A stalk was not observed. Eleven cells were found in LM.

### Remarks

Observed organism was frequently found in samples from the Black Sea and White Sea (unpublished). We can not exclude the possibility of investigating a dividing or dying cells of some known species. Very thin obscure flagellum also known for flagellate species *Ministeria vibrans* (Mylnikov et al. 2019). Presence of longitudinal ventral groove is typical for Jakobida Cavalier-Smith, 1993. Among Jakobida there are some species in the family Histionina Cavalier-Smith, 2013, which also attaches to the substrate and has a very similar cell shape and size: *Histiona aroides* Pasher, 1942, *H. velifera* (Voigt, 1901) Pasher, 1943, *Reclinomonas americana* Flavin et Nerad, 1993, and *R. campanulata* (Penard, 1921) Flavin et Nerad, 1993. These species, however, differ by their possession of lorica and clearly observable flagella. Aloricate Jakobids (such as *Ophirina amphinema* Yabuki et al., 2018, *Andalucia godoyi* Lara et al., 2006, *Moramonas marocensis* Strassert et al., 2016, etc.) are characterized by oval or bean-shaped cells with thick visible flagella, as well as typically swimming cells, not attached to the substrate (Lara et al. 2006; Strassert et al. 2016; Yabuki et al. 2018).

**“Protist 3”** (Fig. 7a–e) – observed in sample point 3c.

**Fig. 7.**
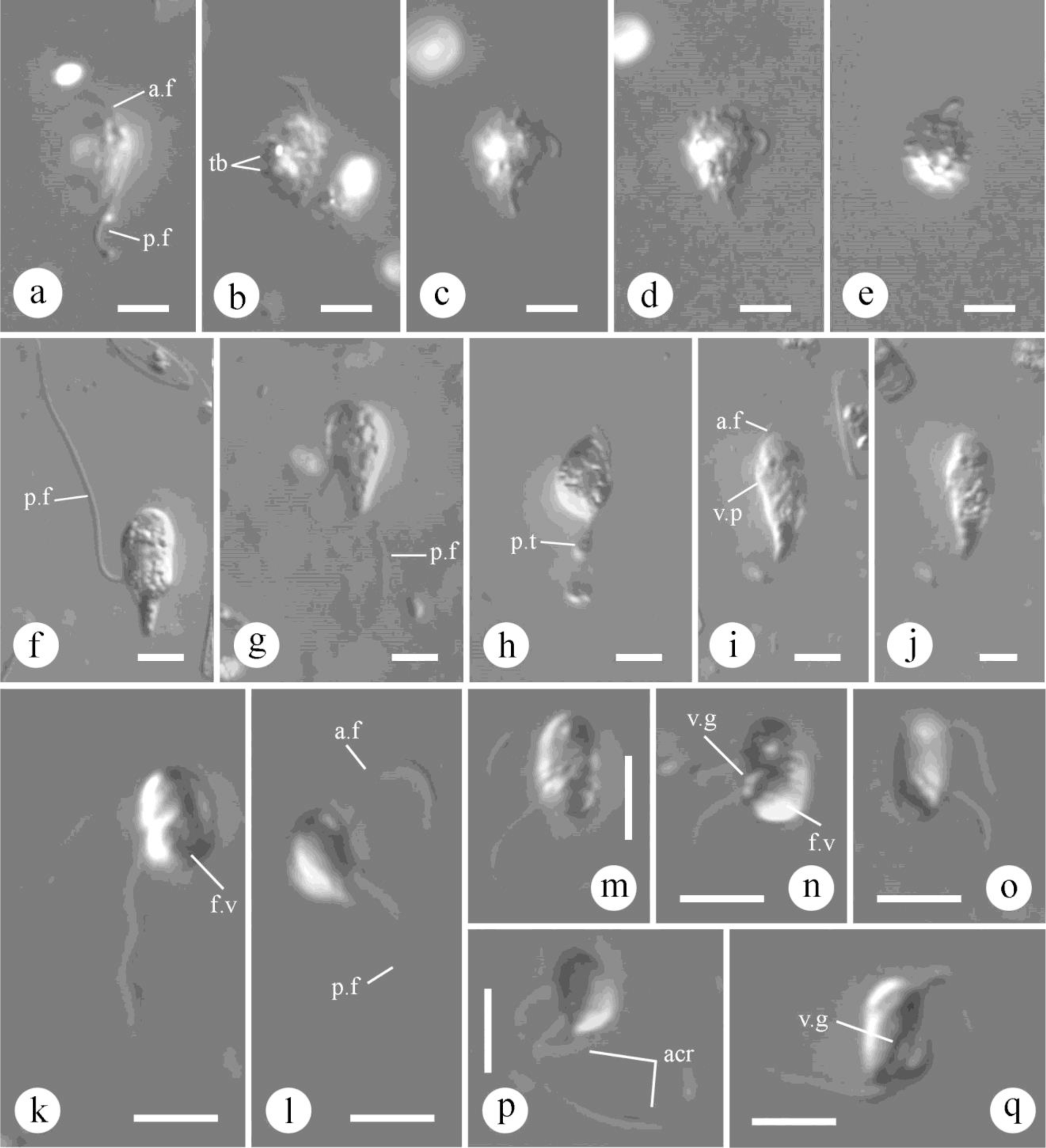
Morphology of observed heterotrophic flagellates (DIC): a–e – “Protist 3”; f–j – “Protist 4” (f – non-swimming cell; g–j – swimming cells); k–q “Protist 5” (l, p – narrow starved cells; k, m, n – saturated cells with prominent food vacuoles; n–q – ventral groove (depression)). Abbreviations: acr – acronema; a.f – anterior flagellum; f.v – food vacuole; p.f – posterior flagellum; p.t – protoplasmic tail; tb – tubercles; v.g – ventral groove; v.p – ventral protrusion. Scale bar: 5 µm.

### Observations

Cell oval in outline, rigid, with rounded anterior end, narrowed posterior end, and convex dorsal side. Not metabolic. Cell length is 7.5–10.0 μm, cell width is 5.5–6.5 μm.

The entire surface of the cell is covered with prominent tubercles. Flagella thick, non-acronematic, and insert apically from the anterior end of the cell. Anterior flagellum is 6.5–8.0 μm in length, directed anteriorly, slightly curved to the dorsal side (forming an arc), and moving very little.

Posterior flagellum is 14–17 μm in length, trailed behind the cell. Flagellates slowly glide or swim. Swimming cells vibrates together with the flagella, slowly rotates in different directions, with flagella keep the position like in gliding cells. Eight cells were examined in LM.

### Remarks

Observed organism most similar to glissomonads (*Allapsa*, *Sandona*, *Heteromita* etc.) and ancyromonads (*Ancyromonas*, *Planomonas*) in shape and size of the cell. However, these species possess less rigid cells, and a smooth cell surface without tubercles. Also, none of these species have a fixed arc position of flagella.

**“Protist 4”** (Fig. 7f–j) – observed in sample points 2d, 6.

### Observations

Cell is 19.5–28.0 μm in length, 8–11 μm in width in the expanded part. Cells pear-shaped, highly metabolic, with posterior protoplasmic tail. Flagella inserts subapically from ventral protrusion. Anterior flagellum thin, slightly noticeable during cell motility, directed forward and undulating. Posterior flagellum thicker, vibrates with a constant amplitude of approximately 10 μm. Cells move slowly and swim in a straight line near the substrate, with a little trembling. Feeding behavior was not recorded, but probably food particles are captured by the caudal or ventral pseudopodia. Specimens were noted both in hypersaline and marine biotopes, with no difference in morphology. Six cells were examined in LM.

### Remarks

Similar cell shape and trembling movement characterized for flagellate genera *Aquavolon*, *Tremula*, and *Lapot*. However, these genera characterized by a clear diagnostic feature: the basal part of the posterior flagellum passes inside the cell and emerge at the central part of the body from the ventral opening. Species of the genus *Tremula* (*T. longifila* Howe et Cavalier-Smith, 2011 and *T. vibrans* (Sandon, 1927) Cavalier-Smith, 2011) glide on the substrate (Howe et al. 2011), while the studied organism was only observed swimming. Species of the genus *Aquavolon* (*A. dientrani* Tikhonenkov, Mylnikov et Bass, 2018 and *A. hoantrani* Tikhonenkov, Mylnikov et Bass, 2018) constantly rotate around the longitudinal axis of the cell during swimming (Bass et al. 2018), while the studied organism swam straight. A similar manner of movement has been described for the species *Lapot gusevi* Tikhonenkov, Mylnikov, Irwin et Keeling, 2019 (Irwin et al. 2019), however, the latter is distinguished by a flattened cell and the presence of few short wide pseudopodia extending from the entire surface of the cell. In addition, known species of *Lapot* and *Aquavolon* were recorded only from freshwater.

**“Protist 5”** (Fig. 7k–q) – observed in sample point 2a.

Observations: Small and very fast moving oval cells, 3.5–7.0 μm in length, 2.5–4.5 μm in width, with two acronematic flagella and anterior nucleus. Flagella inserts from the ventral groove (depression) which begins just above the middle of the cell and goes obliquely to the posterior end of the cell. Posterior flagellum longer than the anterior one, and both are longer than the cell. Flagella and their beating are almost invisible when the cell is moving. Cells swim rapidly and chaotically, always rotate in different planes, and change the direction of movement frequently. Sated cells are wide-oval with prominent posterior food vacuole (Fig. 7k, m, n). Starved cells are narrower, one side is more flattened and other is more roundish, with pointed posterior end (Fig. 7l, p). Eight cells were examined in LM.

### Remark

Presence of large posterior food vacuole may indicate a eukaryotrophic feeding mode of this protist. Fast moving cells with ventral groove resemble colponemids, *Acavomonas,* and *Ancoracysta* (Janouškovec et al. 2017; Tikhonenkov et al. 2014). But the latter are twice the size and slower moving cells. Also, longitudinal grooves in colponemids and *Ancoracysta* do not go obliquely. Starved cells resemble some jakobids and *Cafeteria*-like stramenopiles in body plan, but their movement and behavior is absolutely different. Cells with planar ventral and convex dorsal surface are similar to bacteriovorous *Malawimonas* and some *Carpediemonas*-like organisms. But their flagella insert more anteriorly and are well defined. Also, *Malawimonas* swims in a straight line and known only from freshwater (O’Kelly and Nerad 1993).

## DISCUSSION

The morphology of many of the observed species was slightly different from previous descriptions of any protist. Seven flagellate species were identified only to a genus level: *Thecamonas* sp., *Colpodella* sp., *Cyranomonas* sp., *Goniomonas* sp., *Petalomonas* sp., *Ploeotia* sp. 1, and *Ploeotia* sp. 2. These organisms may represent not yet described species of these genera, but further studies are needed. Among the 86 heterotrophic flagellates and 3 centrohelids encountered in this survey, five heterotrophic flagellates (“Protist 1”, “Protist 2”, “Protist 3”, “Protist 4”, and “Protist 5”) and one сentrohelid heliozoan (*Heterophrys*-like organism) were not identified even to the genus level. Of them, several of the flagellate protists have a unique morphology and may represent undescribed lineages of even a high taxonomic level. From the morphological perspective, we speculate that “Protist 1”, “Protist 3”, and “Protist 4” may be unknown lineages of marine cercozoans. “Protist 5” (resembling *Ancoracysta* and colponemids) and “Protist 2”, which shows no significant similarity to known unicellular eukaryotes, may represent unknown (perhaps deep) branches of the tree of eukaryotes. Several such “weird- looking” protists have indeed recently been shown to represent new lineages (Janouškovec et al. 2017; Eglit, Simpson 2018; Eglit et al. 2017). The under-studied and puzzling *Hemimastix* similarly turned out to be a new, high-ranking phylogenetic lineage of eukaryotes (Lax et al. 2018). Predatory protists with Colponemid-like morphology and behavior are also very intriguing and evolutionarily important, as they are falling in severaldifferent parts of the tree, and contribute to the understanding of the origin of photosynthesis, parasitism, and evolution of mitochondrial genome (Gawryluk et al. 2019; Janouškovec et al. 2013, 2017; Tikhonenkov et al. 2020).

The greatest number of identified flagellate species seem to be members of the SAR supergroup (29 species), Excavates (27), and Obazoa (17). Lower number of flagellate species are apparently members of Cryptista (3), “CRuMs” (2), and Ancyromonadida (1). Six species are of the uncertain systematic position. Centrohelids fall into three families: Raphydocystidae (1 species), Choanocystidae (1), and Heterophryidae (1). All described protist species are new for marine waters of Curaçao, as well as for the Caribbean Sea.

More than half of the identified species of heterotrophic flagellates (46 species, 63%) and all identified centrohelid heliozoans were previously recorded not only from marine, but also from freshwater habitats and can be considered as euryhaline. Twenty-seven flagellate species (37%) were previously reported only from marine and saline inland waters: *Acanthocorbis camarensis*, *Acanthoeca spectabilis*, *Bicosoeca maris*, *Caecitellus parvulus*, *Clautriaviabiflagellata*, *Colponema marisrubri*, *Cyranomonas australis*, *Diplonema ambulator*, *Discocelis punctata*, *Clissandrainnuerende*, *Goniomonas pacifica*, *Halocafeteria seosinensis*, *Mantamonas plastica*, *Massisteria marina*, *Metromonasgrandis*, *Paraphysomonas foraminifera*, *Percolomonas denhami*, *P. similis*, *Petalomonascantuscygni*, *Ploeotia punctata*, *P. robusta*, *Protaspategere*, *Salpingoeca infusionum*, *Stephanoeca cupula*, *S, diplocostata*, *S. supracostata*, *Volknus costatus*.

The most frequent flagellate species in investigated sampling points were *Ancyromonas sigmoides*, *Cafeteria roenbergensis*, *Goniomonas truncata*, *Lentomonas azurina*, *Metopion fluens*, *Metromonas grandis*, *Neobodo designis*, *Petalomonas poosilla*, *Pseudophyllomitus apiculatus*, and *Rhynchomonas nasuta*; they were found in more than 20% of observed samples. Thirty flagellates were rare and found only in one sample. Each of the three observed centrohelid species was observed at only one sampling site.

The average number of species of flagellates and heliozoans in the studied samples is relatively low, 7.0 and 0.1 respectively. Studied marine biotopes have a significantly greater species diversity of flagellate species (total of 83 species, 7.2 on average), compared with hypersaline biotopes, (total of 17 species, 5.0 on average). Common for both marine and hypersaline waters were 15 flagellates: *Amastigamonas debruynei*, *Ancyromonas sigmoides*, *Bicosoeca maris*, *Caecitellusparvula*, *Ciliophrys infusionum*, *Metopion fluens*, *Ministeria vibrans*, *Neobodo designis*, *N. saliens*, *Petalomonas poosilla*, *Ploeotia oblonga*, *Ploeotia* sp. 2, *Podomonas griebenis*, *Rhynchomonas nasuta*, and *Rhynchopus amitus*. *Halocafeteria seosinensis* and *Goniomonas* sp. were found only in hypersaline habitats.

Among observed types of biotope, most flagellate species were found on surface of corals (total of 57 species, on average – 8.1 species per sample) and surface of sponges (39 and 7.2 respectively). These biotopes also were more similar in their species composition compare to other types of habitats (Fig. 8). But the largest number of species in the sample on average was found in coral sand (34 and 13.7). Lower total and average number of flagellate species found in sand samples (24 and 3.9), water column (18 and 6.3), clay (8 and 4.5), and *Sargassum* algae wrings (2 and 2.0). Centrohelid heliozoans were observed only in scraping from sponge (*Raphydocystis bruni*), sand (*Heterophrys*-like organism), and water column (*Choanocystis perpusilla*).

**Fig. 8.**
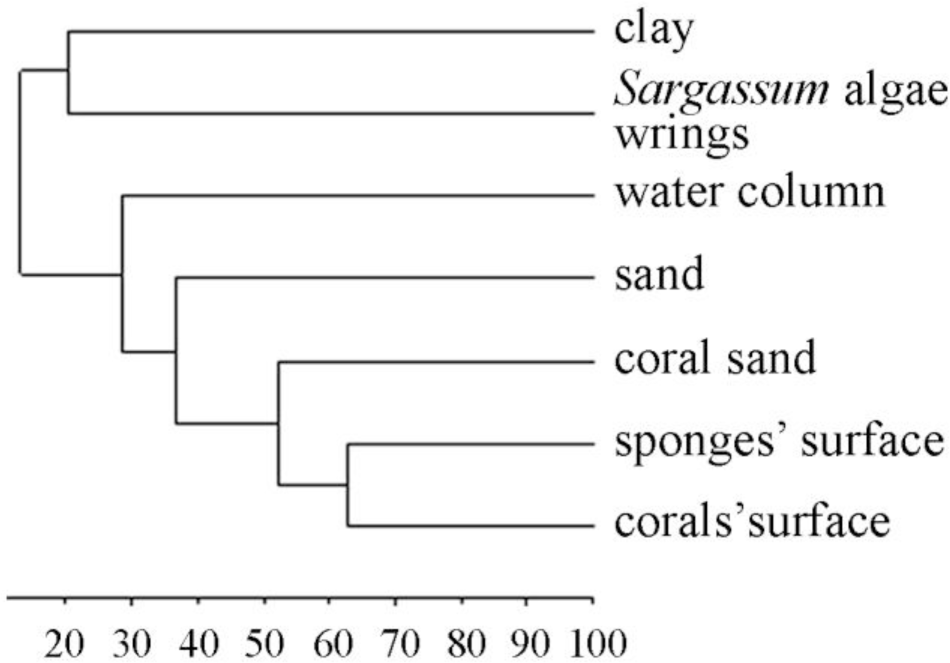
Dendrogram showing the Dice similarity (%) of studied types of biotopes by species composition of heterotrophic flagellates.

The vast majority (89%) of taxa identified to the species level are characterized by a generally wide geographical distribution, and have previously been recorded from all hemispheres (North, South, West, and East), as well as from equatorial, tropical or subtropical, temperate, and polar regions. These (morpho)species can be considered cosmopolites (marked with “c”). Seven species were previously described only from two regions: *Colponema marisrubri*, Red Sea (Tikhonenkov 2009) and Black Sea (Prokina et al. 2018); *Halocafeteria seosinensis*, Korea (Park et al. 2006) and South of European part of Russia (Plotnikov et al. 2011; Prokina 2020); *Petalomonas cantuscygni*, Brazil (Larsen and Patterson 1990) and U.K. (Cann and Pennick 1986); *Ploeotia punctata*, Australia (Al-Qassab et al. 2002; Larsen and Patterson 1990; Patterson and Simpson 1996) and Hawaii (Larsen and Patterson 1990); *Ploeotia robusta*, Australia (Al-Qassab et al. 2002; Patterson and Simpson 1996) and Hawaii (Larsen and Patterson 1990); *Mantamonas plastica*, U.K. (Glücksman et al. 2011) and Australia (Lee 2015); “*Heterochromonas opaca*”, Australia (Lee and Patterson 2000) and Korea (Lee 2002). Among the listed rare species, about half (3) are relatively recently described (after 2000), and further surveys may expand data on their distribution. One species, *Clautriavia biflagellata*, was previously described only once, from Canada (Chantangsi and Leander 2010).

Among identified heliozoans, *Choanocystis perpusilla* was recorded from all hemispheres and all above-listed climatic regions. *Raphydocystis bruni* also has been observed in allhemispheres but only in temperate and tropical regions.

Nevertheless, the conclusions on geographical distribution of protists are highly dependent on which species concept is applied (Azovsky et al. 2016). Here we followed the “morphospecies concept”. However, analysis of the ribosomal RNA genes of several common flagellate morphospecies have shown that they can be represented by morphologically indistinguishable but genetically different strains (von der Heyden and Cavalier-Smith 2005; Koch and Ekelund 2005; Pfandl et al. 2009; Scheckenbach et al. 2006; Scoble and Cavalier-Smith 2014a). Some of these strains appeared to be cosmopolitan, but others were not. In contrast, some benthic cercomonads, the stramenopile *Cantina*, the craspedid choanoflagellate *Codosiga,* and several acanthoecid species have all been shown to have no detected genetic divergence even between geographically distant populations (Bass et al. 2007; Nitsche and Arndt 2015; Stoupin et al. 2012; Yubuki et al. 2015).

Many regions of the world remain insufficiently studied for flagellates and even more so for heliozoans. Indeed, under-sampling is currently the key factor affecting our understanding of protists diversity and distribution (Azovsky et al. 2020). To illustrate this, the species accumulation curve for studied Curacao samples (Fig. 9) fit well (*R*^2^=0.98) with power function *S*=9.36*N*^0.63^. The curve does not flatten out (power coefficient is more than 0.5), so the species list obtained for these sites in Curacao is far from being complete, and each new sample should yield new species.

**Fig. 9.**
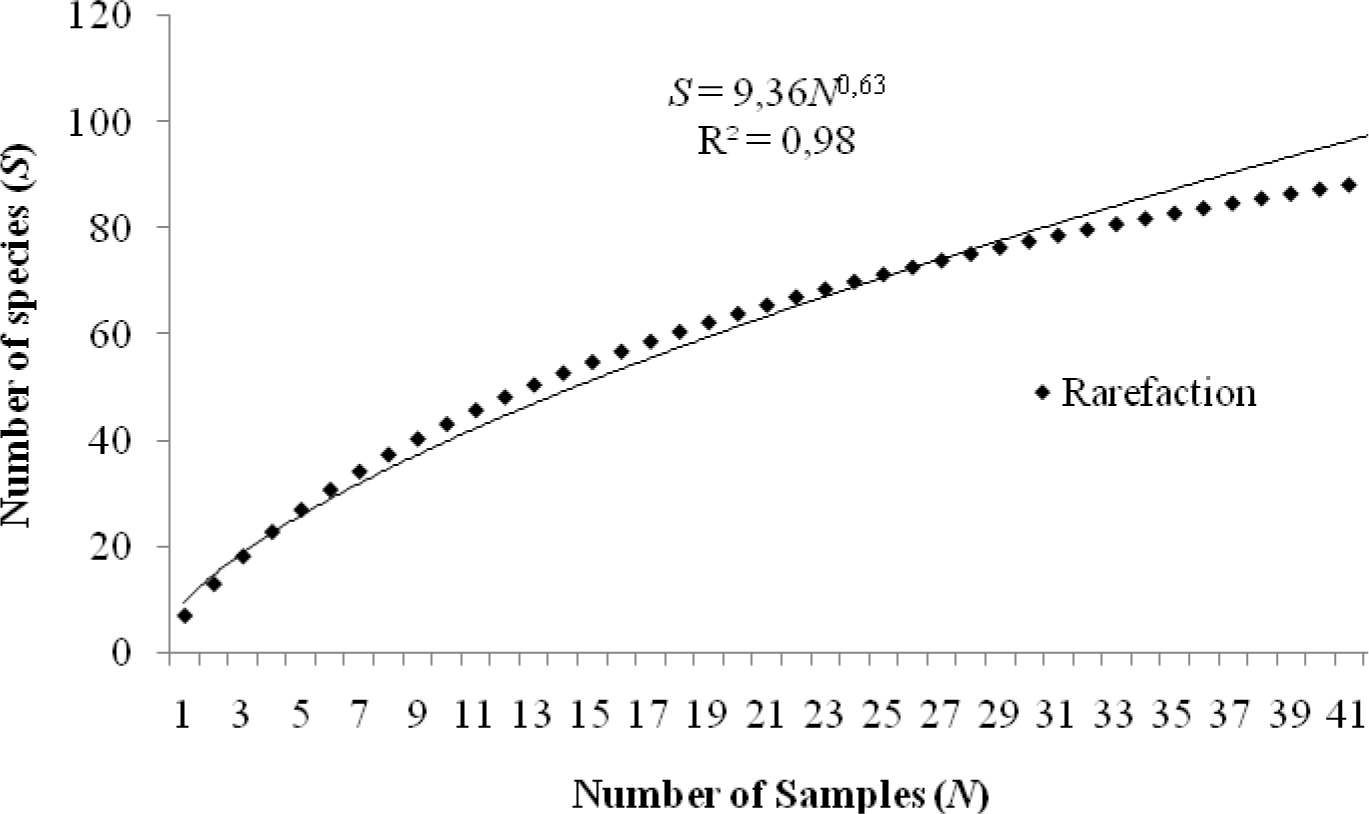
Species accumulation curve.

Undoubtedly, there are many species unaccounted for in each protist diversity survey, and there are many unknown new species in natural ecosystems. Their discovery and description will not only clarify our understanding of distribution patterns of microeukaryotes, but may also contribute to resolving previously puzzling evolutionary and ecological problems. Our observations from Curacao reveal many obscure species identified as “sp.” or not identified at all.

Future work on these protists and molecular investigation in clonal cultures is very promising. Successful isolation in culture largely depends on an understanding of their biology and feeding mode, which can be clarified through observation in natural samples. Alternatively, single-cell transcriptomics is a very efficient approach to study molecular diversity, biology, and the functional state of even small sized protists (Gavelis et al. 2015; Ku and Sebé-Pedrós 2019; Liu et al. 2017; Onsbring et al. 2019), which can be used for ‘uncultivable’ cells. Future discoveries and investigations of flagellates observed but unidentified here, and other unusual protists, will complement the known diversity and biogeography of microbial eukaryotes and can also advance evolutionary research where these organisms represent new branches of the eukaryotic tree.

## Author contributions

Light and electron microscopy, species identification, preparation of species descriptions and illustrations, writing of the original draft (KIP, DVT); statistical analysis (DVT); fieldwork and supervision (DVT, PJK); funding acquisition (DVT, PJK); manuscript review and editing (PJK).

## Acknowledgments

This work was supported by grant from the Russian Foundation for Basic Research (grant no. 20-34-70049) and carried out within the framework of the project no. АААА-А18-118012690098-5 of the Ministry of Education and Science of the Russian Federation. We thank The Gordon and Betty Moore Foundation for travel support, Mark Vermeij and CARMABI research station for field sampling support, and Emma George for help with sample collection.

---

**Table.1.** Characteristics of sampling points

| Habitat | Coordinates, N,<br>W | Date | Biotope | Salinity, ‰ | Depth, m |
| --- | --- | --- | --- | --- | --- |
| 1. Turtle lagoon BokaAscension | 12°16'22.0"<br>69°03'14.0" | April 20,<br>2018 | 1a. Interstitial<br>sand water<br>1b. Sand | 35 | 0 |
| 2. The eastern point of<br>the Curacao island | 12°12'32.3"<br>68°48'58.8" | April 24,<br>2018 | 2a. Sponge<br>2b. Coral<br>2c. Coral sand<br>2d. Sand<br>2e. *Sargassum*<br>algae wrings | 35 | 24.7 |
| 3. Bay SantaMartha | 12°16'05.1"<br>69°07'43.7" | April 26,<br>2018 | 3a. Sponge<br>3b. Coral<br>3c. Sand<br>3d. Coral sand | 35<br>35 | 20<br>0 |
| 4. "Water Factory"<br>sampling site near water<br>treatment plant | 12°06'34.0"<br>68°57'14.9" | April 19,<br>2018 | 4a. Sponge<br>4b. Coral<br>4c. Sand<br>4d. Water<br>column | 35 | 12–18 |
| 5. The western point of<br>the Curacao island | 12°22'31.7"<br>69°09'29.7" | April 26,<br>2018 | 5a. Sponge<br>5b. Coral<br>5c. Sand | 35 | 20 |
| 6. Reef near the Carmabi<br>research station | 12°07'19.8"<br>68°58'09.4" | April 22,<br>2018 | 6. Coral | 35 | 20 |
| 7. Hilton Hotel Beach | 12°07'17.3"<br>68°58'08.8" | April 24,<br>2018 | 7. Coral sand | 35 | 0 |
| 8. Hypersaline lagoon<br>Saliña Sint Maria | 12°12'45.7"<br>69°03'15.3" | April 20,<br>2018 | 8a. Black clay<br>8b. Water<br>column | 52 | 0 |

